# Human TIMELESS, a potential circadian clock regulator, plays an essential role in survival and reawakening of metastasis-initiating cells in bone

**DOI:** 10.1101/2025.03.18.643632

**Authors:** Hirokazu Shimizu, Lei Wang, Masumi Tsuda, Reo Maruyama, Yusuke Saito, Kohei Kumegawa, Masato Takahashi, Rie Horii, Hiroaki Suzuki, Kenichi Watanabe, Ryuta Arai, Masaaki Murakami, Norimasa Iwasaki, Shinya Tanaka

**Affiliations:** Department of Orthopaedic Surgery, Faculty of Medicine and Graduate School of Medicine, Hokkaido University, Sapporo, Japan; Department of Cancer Pathology, Faculty of Medicine, Hokkaido University, Sapporo, Japan; Department of Musculoskeletal Oncology, NHO Hokkaido Cancer Center, Sapporo, Japan; Institute for Chemical Reaction Design and Discovery (WPI-ICReDD), Hokkaido University, Sapporo, Japan; Project for Cancer Epigenomics, Cancer Institute, Japanese Foundation for Cancer Research, Tokyo, Japan; Cancer Cell Diversity Project, NEXT-Ganken Program, Japanese Foundation for Cancer Research, Tokyo, Japan; Department of Breast Surgery, Faculty of Medicine and Graduate School of Medicine, Hokkaido University, Sapporo, Japan; Department of Pathology, Saitama Cancer Center, Saitama, Japan; Department of Pathology, NHO Hokkaido Cancer Center, Sapporo, Japan; Department of Breast Oncology, NHO Hokkaido Cancer Center, Sapporo, Japan; Department of Orthopaedic Surgery, Aiiku Hospital, Sapporo, Japan; Department of Surgical Pathology, Hokkaido University Hospital, Sapporo, Japan; Division of Molecular Psychoneuroimmunology, Institute for Genetic Medicine and Graduate School of Medicine, Hokkaido University, Sapporo, Japan; Quantum Immunology Team, Institute for Quantum Life Science, National Institutes for Quantum Science and Technology (QST), Chiba, Japan; Division of Molecular Neuroimmunology, National Institute for Physiological Sciences, National Institutes of Natural Sciences, Okazaki, Japan; Institute for Vaccine Research and Development (HU-IVReD), Hokkaido University, Sapporo, Japan

**Keywords:** TIMELESS, circadian clock regulator, bone metastasis, metastatic latency

## Abstract

Bone metastasis is becoming increasingly common globally. In such cases, cancer cells disseminate from the primary site to bone, subsequently entering a dormant state. Following a certain incubation period, the cells reawaken and grow, forming a metastatic mass. These dormant cells are called metastasis-initiating cells (MICs), but their survival and reawakening are poorly understood. Here, we established an *in vivo* MIC-reawakening mouse model in bone. Microarray analysis demonstrated enrichment for mitochondrial oxidative phosphorylation (OXPHOS) and fatty acid synthesis in isolated MICs. In addition, transcriptional regulator TIMELESS was identified as an independent poor prognostic factor by using clinical transcriptomic datasets, which was validated by immunostaining on 209 breast cancer cases. TIMELESS-deficient breast, prostate, and bladder cancer cell lines exhibited reduced viability and MIC reawakening in bone. Single-cell analysis on the epigenetic landscape of MICs revealed that motif activities of CLOCK and BMAL1/ARNTL were enhanced in a cluster with increased TIMELESS accessibility scores. As TIMELESS regulates stemness through the OXPHOS metabolic state, an inhibitor targeting the metabolic pathway, MP-A08, was identified to suppress MIC reawakening and growth in bone more efficiently than cisplatin. These results suggest that TIMELESS regulates survival and reawakening in MICs and MP-A08 could contribute to adjuvant therapeutic strategies.

## Introduction

In the process of cancer metastasis, cancer cells derived from a primary lesion reach and reside at a secondary site in a dormant state^1^. A small proportion of these resident cells reawaken at a certain timepoint and proliferate to form a large mass^1^. These dormant cancer cells can subsequently form macro-metastases and are called to be metastasis-initiating cells (MICs)^1,2^. Upon reaching distant organs including bone, MICs might remain in a latent and non-proliferative state for several months to years. Latent MICs can evade immune surveillance by natural killer (NK)/T cells by downregulating the expression of NK cell-activating ligands and major histocompatibility complex (MHC) class I molecules^3,4^. In addition, stromal TGF-β and BMP signals in distant organs inhibit the proliferation of MICs^5,6^ and intratumoral stimulator of interferon gene activity also suppresses metastatic outbreak^7^. These inhibitory mechanisms may contribute to the dormancy of MICs, which is clinically known as metastatic latency^3,8,9^.

The cancer metastatic latency of MICs can be divided into three stages: dissemination to distant organs, transition to quiescence, and awakening from the dormancy^1,2^. Although latent MICs and their reawakening are a major concern in a clinical context, little is known about the cancer-intrinsic trigger for the survival and reawakening of MICs^2,10^. Thus, identification of the positive regulators of these processes could lead to the development of therapeutic agents for MICs, which could in turn facilitate the establishment of potentially curative cancer treatments.

Cancer cells dynamically adapt their metabolism during each step of metastasis^11^. Generally, cancer cells metabolically generate energy using glycolysis, rather than mitochondria, via the Warburg effect. However, in metastatic cancer cells, the metabolic program switches to enhance mitochondrial oxidative phosphorylation (OXPHOS) in bone metastasis^11^. The machinery involved in the metabolic adaptation of metastatic cancer cells is controlled by transcriptional regulation^12^, but the majority of metabolic studies have focused on the early stage of metastasis, including the migration of primary cancer cells and their invasion into blood vessels^13^, rather than the reawakening of MICs at metastatic sites.

The dynamics of metastatic progression has been reported to be closely related to circadian rhythm. The intravasation of circulating tumor cells (CTCs) occurs during sleep in breast cancer patients, and single-cell sequencing of CTCs revealed marked upregulation of mitotic genes at night in humans^14^. The coupling of circadian clock and cell cycle in a physiological state results in timed mitosis and rhythmic DNA replication^15^. Furthermore, disruption of the circadian clock causes accelerated cancer onset and progression^16,17^. Because latent MICs have low mitotic capacity, the mechanisms of survival and reawakening of MICs may be related to circadian rhythms.

With reference to the experimental latency model in bone^18–20^ based on tumor mass dormancy^18^, we here created a novel model of bone metastasis that accurately recapitulates the MIC reawakening system, and identified that the circadian rhythm-related transcriptional regulator TIMELESS plays a crucial role in the reawakening of MICs. As such, we show that inhibitors of the TIMELESS-related pathway may be potent therapeutic agents to prevent cancer metastasis.

## Results

### Establishment of an *in vivo* MIC-reawakening model for bone metastasis of breast and bladder cancers

To examine the mechanism of MIC reawakening in bone metastasis, we employed an *in vivo* MIC-reawakening model in which cancer cells were injected via the tail artery of mice^21^. Bioluminescence imaging (BLI) demonstrated that 1.0 × 10^6^ cells of human breast cancer MCF7 and bladder cancer UM-UC-3 cell lines injected into mice showed a certain signal intensity at the knees on day 3 (Fig. 1a and 1b). These models exhibited a latent period of approximately 14 days, as previously reported^18–20^. Note that we carefully ruled out the possibility of failure of the detection sensitivity within 14 days, since the measured intensity of bioluminescence was directly proportional to the number of subcutaneously implanted cells ranged from 10 to 320 (data not shown). It was consistent with a previous report^22^. MICs subsequently awakened and started to regrow (Fig. 1a and 1b) up to day 42 (Fig. 1a). On day 14 of the latent period, a single tdTomato-labeled UM-UC-3 MIC could be detected in the surrounding knee trabecular structure of femoral bone by fluorescence microscopy (Fig. 1c and 1d).

**Figure 1:**
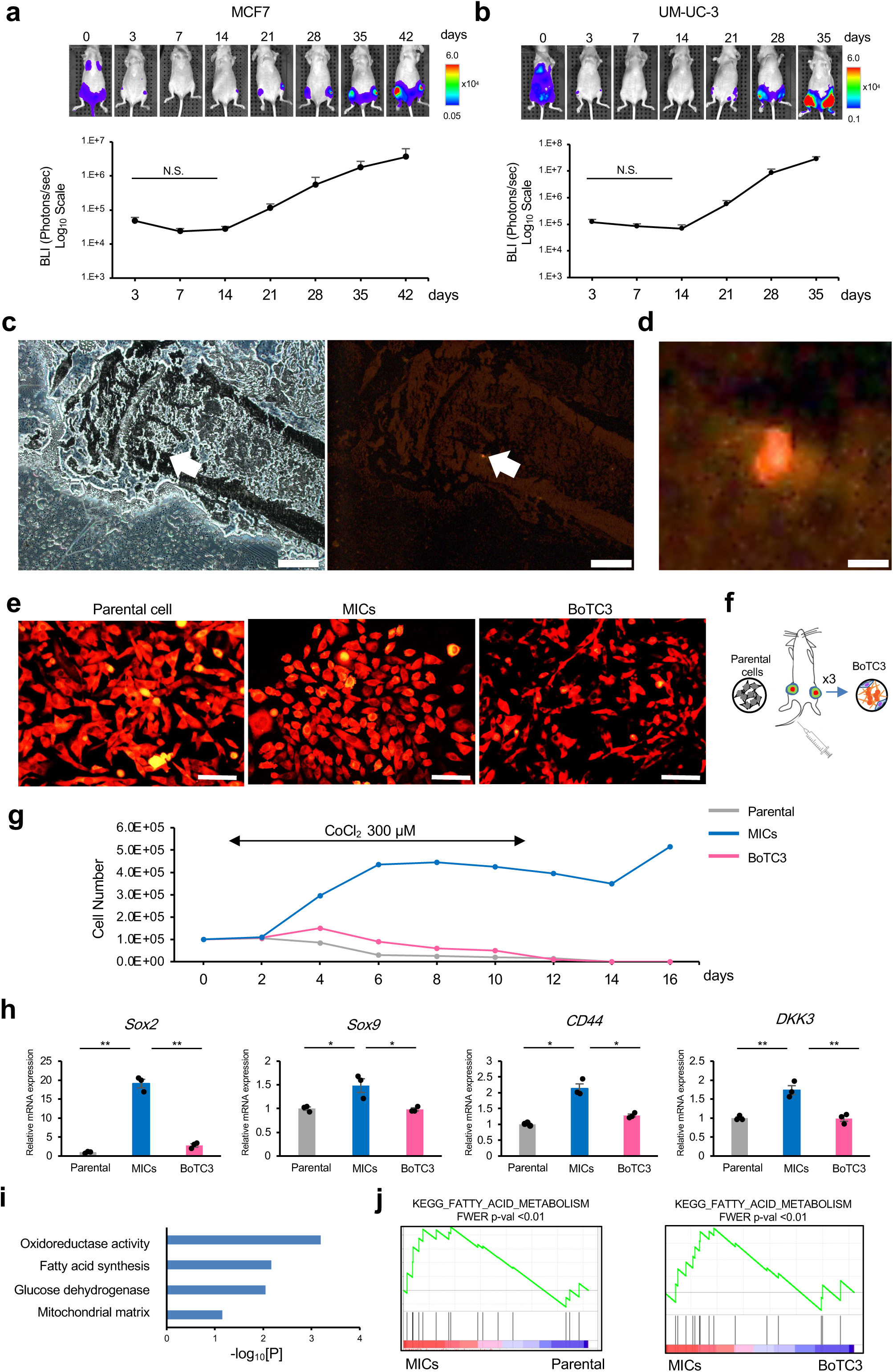
*In vivo* MIC-reawakening models and profiles of isolated MICs. **a-b**, *In vivo* MIC-reawakening models focusing on bone with 14-day latency period and subsequent reawakening using MCF7 (a) and UM-UC-3 (b) cells, (n=6 limbs with metastatic lesion in each cell line) **c**, Representative images of the limb at day 14 after injection by bright-field microscope (left panel) and fluorescence microscope (right panel). Scale bar, 400 µm. **d**, The enlarged image of 1c focusing on the td-tomato positive cell. Scale bar, 20 µm. **e**, Representative images of *in vitro* parental cell, MICs, and BoTC3. **f**, Graphical schematics of establishment of BoTC3. **g**, *In vitro* growth under 300 µM CoCl_2_ condition for parental cells, MICs, and BoTC3. Scale bar, 100 µm **h**, Relative mRNA expression of *Sox2*, *Sox9*, *CD44*, and *DKK3* in parental cells, MICs, and BoTC3. The data represent means ± s.e. (n=3 independent experiments). Two-sided paired t-test were used for statistical analysis. *P* value for comparisons in the graphs are indicated (\**P*<0.05, \*\**P*<0.05) **i**, Gene-expression signatures that associate enrichment in MICs compared with parental cells. Holizontal axis indicate *P* values **j**, Enriched gene sets related to metabolisms in MICs compared with BoTC3 (left) and parental cells (right). MICs: metastasis-initiating cells. BoTC: bone-tropic tumor cell

A single MIC in the latency period may have potential not only to survive in bone but also to reawaken there and form macroscopic metastases. To further analyze the characteristics of MICs, the femoral bone tissues surrounding the knee joint, including bone marrow, of mice at day 14 after UM-UC-3 injection were homogenized. The isolated cells were then cultured *in vitro* for 14 days with G148 to select drug-resistant tumor cells, and the established cells were designated as MICs (Fig. 1e, middle panel). For comparison, mass-forming cells in metastatic bone were isolated on day 35, and these cells were reinjected into the caudal artery, resulting in massive bone metastases. Metastatic cells designated as bone-tropic tumor cells 3 (BoTC3s) were established after repeating this process three times (Fig. 1e and 1f). In contrast to the morphology of the parental cells and MICs, the BoTC3s were spindle-shaped and elongated (Fig. 1e). The BoTC3s also underwent non-proliferative periods^22^ (Extended data Fig. 1a and 1b).

### MICs exhibited metabolic OXPHOS with enhanced fatty acid synthesis

To investigate the *bona fide* character of MICs, the effect of hypoxia, a well-known feature of the stem cell niche, on these cells was assessed by culturing MICs derived from human bladder cancer UM-UC3 cells in culture medium supplemented with cobalt chloride (CoCl_2_), a hypoxia-mimetic agent^23^. It was found that an increase in the number of MICs stopped during CoCl_2_ treatments. After the removal of CoCl_2_, however, the number of MICs began to re-increase (Fig. 1g). In contrast, the viabilities of parental UM-UC-3 or BoTC3 were remarkably decreased under CoCl_2_ treatments (Fig. 1g). These results suggest that MICs enter a quiescent state and can survive under hypoxic conditions that mimic the microenvironment as a cancer stem cell niche that is located in the vicinity of trabecular bone, causing cancer recurrence in patients.

Confirming the identity of the MICs, the established MCF7-derived MICs were found to highly express *SOX2*, *SOX9*, and the *DKK* family of genes (Fig. 1h), as previously reported^3^. Furthermore, MICs exhibited higher levels of WNT5a and its receptor ROR2 together with TRPS1, a related transcription factor (Extended data Fig. 2a), which were reported to play important roles in the attachment of circulating cancer cells to target organs and the suppression of tumor initiation (Extended data Fig. 2a– 2e)^20,24,25^. These results suggest that the isolated MICs are compatible with the metastatic cancer cells in the latent phase. As an analysis of the characteristics of MICs, gene expression profiling by DAVID showed increases in oxidoreductase activity and fatty acid biosynthesis as the most enriched clusters (Fig. 1i). Gene set enrichment analysis (GSEA) also revealed the enriched expression of genes related to fatty acid metabolism and oxidative phosphorylation (OXPHOS) in the MICs compared with the levels in parental cells (Fig. 1j and Extended data Fig. 2f). Focusing on lipid metabolism, enriched pathways in MICs were analyzed using the KEGG database (Extended data Fig. 3a and 3b), which revealed the enrichment of fatty acid synthesis in mitochondria as well as the cytosol. In particular, genes related to the pentose phosphate pathway and to cell cycle checkpoints were enriched in MICs compared with the levels in BoTC3s (Extended data Fig. 2g), whereas steroid synthesis and gluconeogenesis were downregulated in MICs (Extended Data Fig. 2h and 2i). Notably, three genes, namely, *SCD*, *SREBF1*, and CEBPA, which were reported to be crucial factors of lipid metabolism for the growth of brain metastatic breast cancer cells^12^, were downregulated in MICs (Extended Data Fig. 2h).

**Figure 2:**
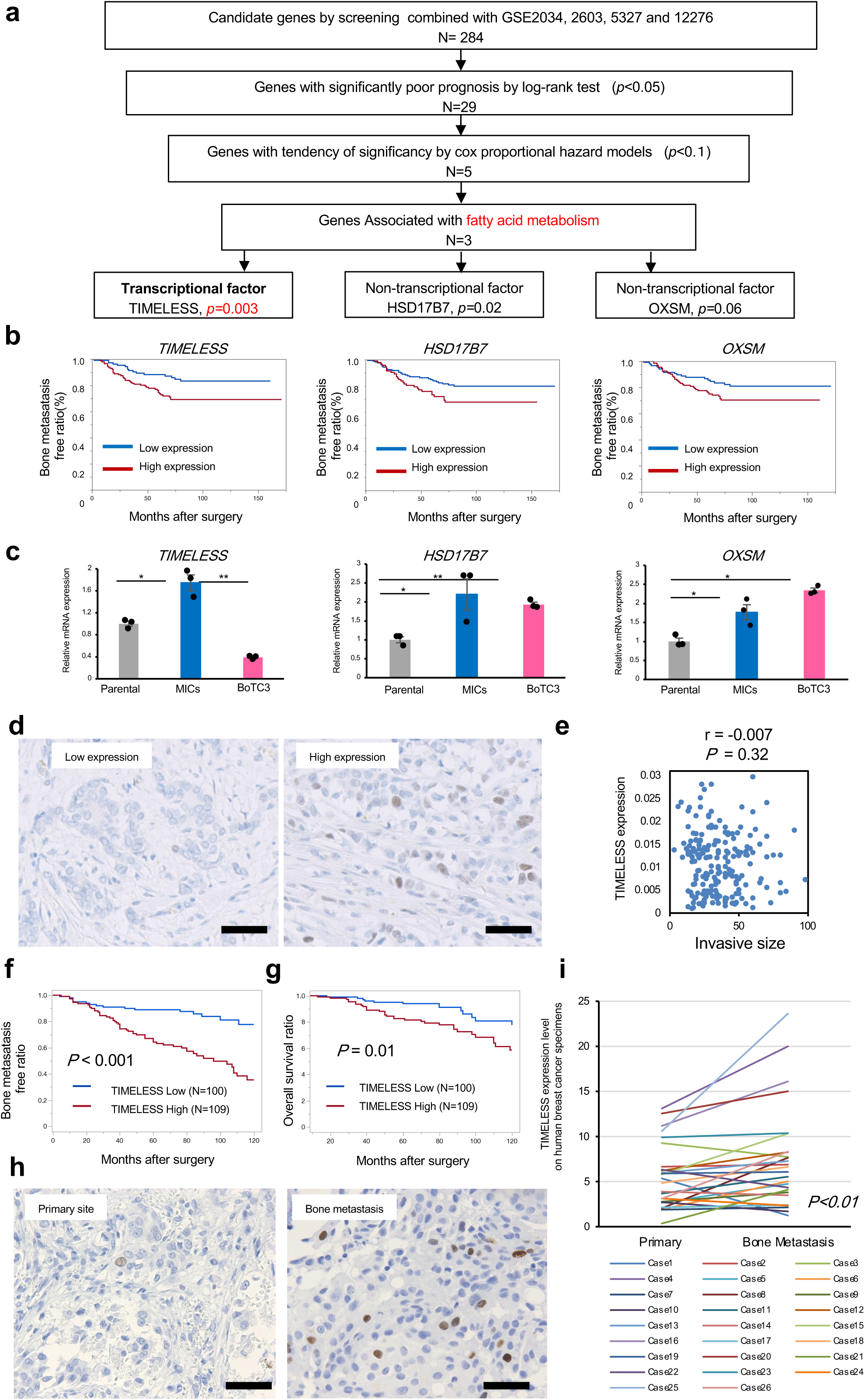
Screening of MIC genes with clinical transcriptome datasets identifies TIMELESS as the independent poor prognostic factor. **a**, The independent poor prognostic genes in upregulated genes in MICs, extracted by a screening model with the MSKCC/EMC cohort primary tumor dataset. **b**, Survival analysis representing the proportion of bone metastasis free patients (from the MSKCC/EMC cohort primary tumor dataset) stratified according to *TIMELESS*, *HSD17B7*, and *OXSM*. Significance was assessed by log ranked test. **c**, Relative mRNA expressions of *TIMELESS*, *HSD17B7*, and *OXSM* in parental cells, MICs and BoTC3. (n=3 independent experiments). **d**, Representative images of TIMELESS staining on ER^+^ stage 2b and 3 breast cancer patient samples. Total sample, n= 209. Scale bar, 200 µm. **e**, Pearson correlation between TIMELESS expression and tumor invasive size in ER^+^ stage 2b and 3 breast cancer patients. r=-0.007, P=0.32. **f**,**g** The proportion of bone metastasis-free patients (**f**) and overall survival analysis of patients (**g**) from the Hokkaido Cancer Center and Saitama Cancer Center cohort primary tumor data set (TIMELESS high, n=109; TIMELESS low, n=100). **h,** Representative images of TIMELESS immunohistochemistry on ER^+^ breast cancer primary and the matched bone metastasis tumor. Total sample, n= 27. Scale bar, 200 µm. **i**, pair match positivity of TIMELESS staining on ER^+^ breast cancer primary and matched bone metastasis tumor (n=26). The data represents means ± s.e.. Two-sided paired t-test were used for statistical analysis. For **f** and **g**, significance was assessed by log ranked test. For **i**, pairs are connected with a line and Wilcoxon signed-rank P value is shown. *P* value for comparisons with the results are indicated (\**P*<0.05, \*\**P*<0.01)

**Figure 3:**
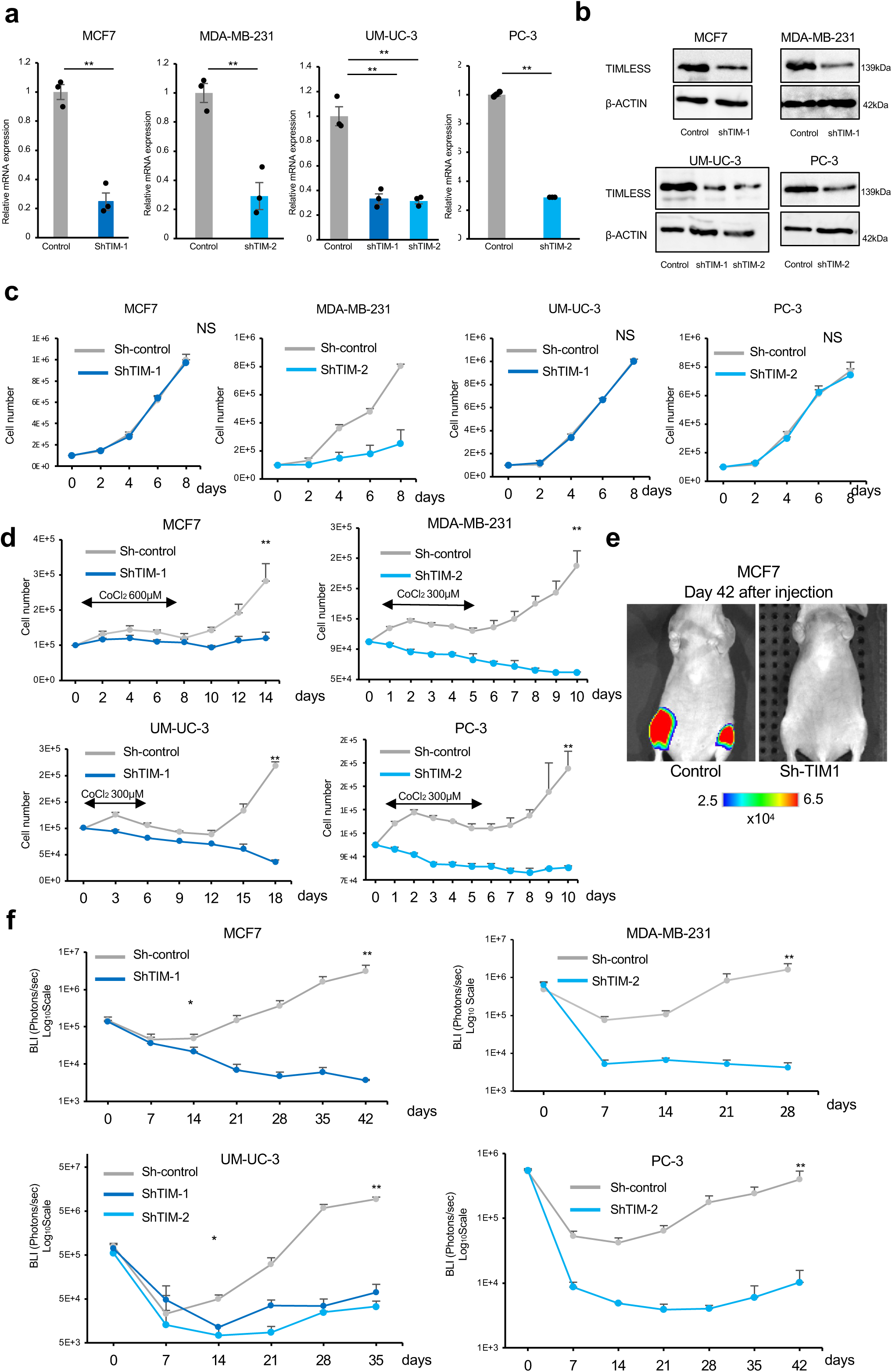
TIMELESS plays an essential role for survival and reawakening of MICs in bone. **a**, Relative mRNA expression in *TIMELESS* knockdown cell lines (shTIM-1 and shTIM-2) in MCF7, MDA-MB-231, UM-UC-3 and PC-3 cells (each n=3 independent experiments). **b**, Western blotting of the TIMELESS knockdown cell lines (shTIM-1 and shTIM-2) in MCF7, MDA-MB-231, UM-UC-3, and PC-3 cells. **c**, *In vitro* cell growth of the TIMELESS knockdown cell lines and controls in MCF7, MDA-MB-231, UM-UC-3, and PC-3 (each n=3). **d**, *In vitro* cell growth with CoCl_2_ condition in the *TIMELESS*-knockdown cell lines in MCF7, MDA-MB-231, UM-UC-3, and PC-3 (each n=5 independent experiments). **e**, Representative images of *in vivo* bone metastasis at day 42 after caudal injection of MCF7 cells (1.0 *×* 10^6^) to mice. **f**, Survival and subsequent metastatic outbreaks of tumor cells in mice receiving caudal arterial injection (MCF7: control (n=8 limbs), shTIM-1 (n=12 limbs); MDA-MB-231: control (n=8 limbs), shTIM-2 (n=8 limbs); UM-UC-3: control (n=8 limbs), shTIM-1 (n=8 limbs), sTIM-2 (n=8 limbs); PC-3: control (n=8 limbs), shTIM-2 (n=8 limbs). For **a**, **c**, **d**, and **f**, significance was assessed by two-sided paired t-tests with the data representing means ± s.e.. *P* value for comparisons with the results are indicated (\**P*<0.05, \*\**P*<0.01). N.S., not significant.

### Screening of MIC genes using clinical transcriptome datasets identified TIMELESS as an independent poor prognostic factor

To identify genes essential for MIC survival and reawakening, microarray analysis was performed, which revealed that 284 genes were upregulated 2.0-fold or more in MICs derived from UM-UC-3 compared with the level in the parental cell line. Screening was performed using the clinical samples of early-stage estrogen receptor-positive breast cancer allocated to a bone metastasis cohort (Fig. 2a; the data are publicly available from Memorial Sloan Kettering Cancer Center/Erasmus Medical Center cohort, GSE 2034 and GSE 2603). Cox proportional hazard models revealed five genes with a p-value of less than 0.10: *TIMELESS* (p = 0.003), *HSD17B7* (p = 0.02), *AP1S1* (p = 0.02), *CDKN3* (p = 0.02), and *OXSM* (p = 0.06) (Fig. 2a and 2b). Among the three genes related to fatty acid metabolism enriched in MICs (Extended Data Fig. 3a and 3b), *TIMELESS* was specifically expressed in MICs derived from UM-UC-3, whereas *HSD17B7* and *OXSM* were upregulated in both MICs and BoTC3 (Fig. 2c). Immunofluorescence of TIMELESS also revealed that the distribution of signal intensity in MICs was bimodal, but unimodal in the parental cells (Extended data Fig. 4a and 4b). The mean signal intensity was significantly higher in MICs than in the parental cells (Extended data Fig. 4c).

**Figure 4:**
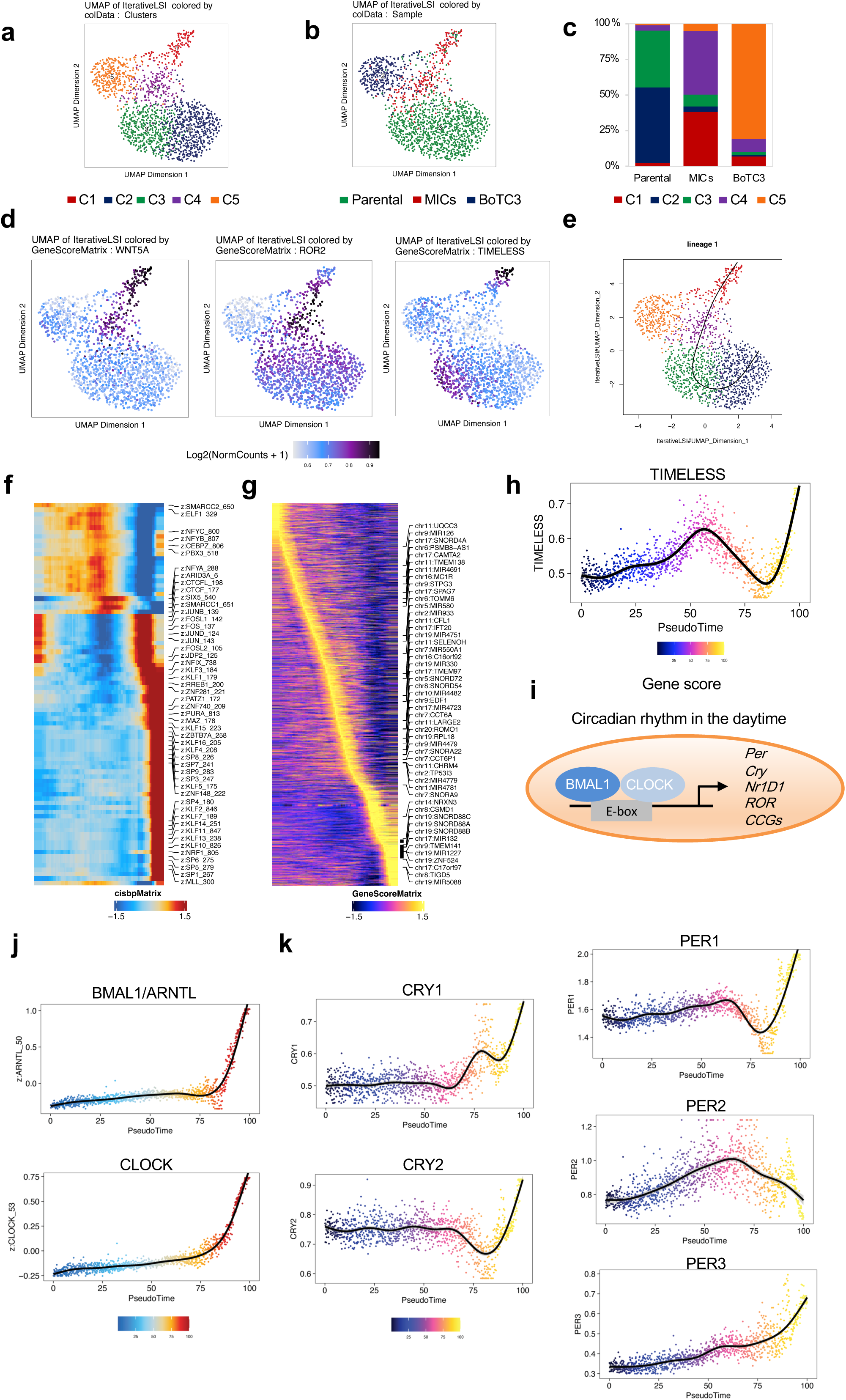
Single-cell epigenomic landscape of MICs shows involvements of core factors for circadian rhythms. **a**, Uniform manifold approximation and projection (UMAP) after iterative latent semantic indexing of scATAC-seq profiles from the parental cells, MICs, BoTC3. Each dot represents a single cell colored by its corresponding cluster. Each cluster number is presented on the UMAP. **b**, The UMAP projection shown in **a** is colored by its corresponding sample. **c**, Ratios of each cluster in the parental cells, MICs, BoTC3 are shown. **d,** UMAP projection colored by gene accessibility scores, reflecting the general chromatin accessibility of the indicated gene. *WNT5a* (left), *ROR2* (middle), and *TIMELESS* (right). **e**, Pseudotime trajectory analysis producing lineage from the cluster 2 to the cluster 1. **f**, Pseudotime heatmap ordering of most variable chromVAR TF motifs across the lineage. **g**, Pseudotime heatmap ordering of gene accessibility scores of the most variable genes across the lineage1. **h**, Gene accessibility scores of TIMELESS along the psuedotime trajectory. **i.** BMAL1/ARNTL and ClOCK increase the transcription of Per and Cry and CCGs during the day. **j,** Motif enrichments of ARNTL and CLOCK along the pseudotime trajectory. **k**, Gene accessibility scores of CCGs along the psuedotime trajectory. CCGs: clock-controlled genes

The screening model was also applied to identify favorable prognostic factors in MICs (Extended Data Fig. 5a–c), with *MGLL* (p = 0.007)*, TXNIP* (p = 0.02), *PRKAR2B* (p=0.04), *G0S2* (p=0.07), and *PORCN* (p = 0.07) being detected. *MGLL* and *TXNIP* are associated with oxidative stress responses^26^. *SNAI2* was detected as a poor prognostic gene using the model with BOTC3s (p = 0.02) (Extended data Fig. 6a and b). A formula for predicting the prognosis was created using the following 11 prognostic factors: *TIMELESS*, *HSD17B7*, *OXSM*, *MGLL*, *TXNIP*, *PRKAR2B*, *G0S2*, *PORCN*, *MIR22*, *TMEM158*, and *SNAI2* (Extended data Fig. 6c and 6d).

To examine the clinical value of *TIMELESS* in predicting the prognosis, surgically resected breast cancer tissues (n=209) were employed for immunohistochemical analysis (Fig. 2d). No correlation was observed between tumor size and expression levels of TIMELESS (Fig. 2e), but the TIMELESS-high group showed early relapse of bone metastases and poor overall survival (Fig. 2f and 2g). In a patient-matched analysis in 26 identical patients with breast cancer, the rate of positivity of TIMELESS immunostaining was significantly higher in bone metastatic tissues than in primary cancer tissues (Fig. 2h and 2i). It should be noted that patient-matched analysis using publicly available transcriptome datasets^27^ validated this (Extended data Fig. 7).

These results suggested that *TIMELESS* was worth investigating in further experiments since it was reported to regulate the production of sphingosine-1 phosphate (S1P) via the Sp1/ACER2/S1P axis^28^, which plays a role in the survival and proliferation of metastatic cancer cells^29^.

### *TIMELESS* knockdown decreased survival of MICs and subsequent reawakening in bone

To examine the role of TIMELESS in MICs, *TIMELESS*-knockdown cells were established using four cell lines of breast, bladder, and prostate cancers; MCF7, MDA-MB-231, UM-UC-3, and PC-3, respectively (Fig. 3a and 3b). Cell growth of MCF7, UM-UC-3, or PC-3 was not affected by *TIMELESS* depletion, whereas the growth of MDA-MB-231 was suppressed (Fig. 3c and Extended data Figure 8a and 8b). Under the hypoxic conditions produced by CoCl_2_ treatment, cell growth was suppressed with the cell number plateauing, and cells re-proliferated after the removal of CoCl_2_ in all four parental cells (Fig. 3d and Extended data Figure 8c). However, in *TIMELESS-* knockdown cells, cell number was decreased during hypoxia, and cell regrowth was not observed even after the hypoxia was released (Fig. 3d and Extended data Fig. 8d).

In the *in vivo* metastatic model, cells metastasized to the knee bones of mice and did not increase in number within 1–2 weeks, designated as the latency period (Fig. 3f). Notably, a significant decrease in cell proliferation by TIMELESS knockdown was observed at day 14 in all four cell lines (Fig. 3f). After 2 weeks, MICs began to grow, representing the reawakening period. *TIMELESS* knockdown resulted in a significant decrease in their survival in femoral bone of mice. At 28 to 42 days after the intra-caudal injection of *TIMELESS*-knockdown cells, marked defects in bone metastasis (44- to 425-fold) were observed (Fig. 3e and 3f). These results clearly demonstrated that TIMELESS plays an essential role in the survival and reawakening of MICs.

### MICs with high chromatin accessibility of TIMELESS and WNT5a had remarkable motif enrichments of CLOCK and BMAL1/ARNTL

As the epigenetic landscape of MICs is largely unknown, chromatin accessibility profiling analysis by droplet-based scATAC-seq was performed on MICs, parental cells, and BoTC3s derived from the bladder cell line. Single-cell-based analysis was conducted since the isolated MICs showed heterogeneity on immunostaining of TIMELESS (Extended data Fig. 8a). The appropriateness of the MICs, parental cells, and BoTC3s in terms of their quality was confirmed^30,31^ (Extended data Fig. 9a–e).

Graph-based clustering was performed based on the chromatin accessibility landscape, resulting in a total of five major clusters (Fig. 4a and 4b). MICs were mainly classified into two clusters (clusters 1 and 4) (Fig. 4c). The gene accessibility score of *Wnt5a* was particularly increased in clusters 1 and 4 (Fig. 4d). Meanwhile, the score of *TIMELESS* was increased in cluster 1, whereas that of *Wnt5a receptor*, *ROR2*, was increased mainly in cluster 4 (Fig. 4d). Finally, the gene accessibility score of *TRPS1*, a transcription factor related to *WNT5a* (Extended data Fig. 2c and 2d), was also enhanced in cluster 4 (Extended data Fig. 10d).

To clarify the mechanisms driving the epigenomic state of MICs, trajectory-based analysis was performed to identify lineages 1 and 2 (Fig. 4e and Extended data Fig. 10a). Dynamic changes in the enrichment of transcription factor (TF)-binding motifs and gene accessibility scores along the trajectory were observed (Fig. 4f and 4g, Extended data Fig. 10b and 10c). In lineage 1, an increase in the gene accessibility score of *TIMELESS* was observed at the late stage (Fig. 4h). Focusing on the feedback loop of mammalian circadian rhythms, *CLOCK* and *BMAL1/ARNTL* increase the transcription of *Period (Per), Cryptochrome (Cry),* and other *clock-controlled genes (CCGs)* during the day (Fig. 4i)^32^. Our results showed remarkable motif enrichments in *CLOCK*, *BMAL1/ARNTL*, and stemness genes at the late stage (Fig. 4j and Extended data Fig. 10f). Gene accessibility scores of *Per*, *Cry*, and other *CCGs* were also increased at the late stage (Fig. 4k and Extended data Fig. 10e). Thus, the motif activities of *CLOCK* and *BMAL1/ARNTL* and gene accessibility scores of *CCGs* were enhanced at the late stage, and the gene accessibility scores of *TIMELESS* and *Wnt5a* were also increased.

### *TIMELESS*-regulated oxidative phosphorylation was crucial for bone metastases

To examine the mechanisms of *TIMELESS*-regulated MIC reawakening, we examined the stemness features and found that the expression levels of stemness-related genes, such as *Sox2*, *Sox9*, and *CD44*, were significantly decreased by *TIMELESS* depletion in PC-3 (Fig. 5a). In addition, in *TIMELESS*-knockdown cells, the levels of *OXSM*, *HSD17B7*, and *HSD17B8*, which are involved in fatty acid synthesis in mitochondria, were significantly reduced in PC-3 (Fig. 5b and 5c and Extended data Fig. 5a). In the sphingolipid salvage pathway, the levels of *alkaline ceramidase 2* (*ACER2*) and *alkaline ceramidase 3* (*ACER3*) were decreased in the *TIMELESS-* knockdown cells derived from PC-3 (Fig. 5d and 5e), resulting in a decrease in the product sphingosine monophosphate (S1P) as confirmed by ELISA (Fig. 5f). In contrast, the expression of S1PR1, the receptor for S1P, was significantly increased (Fig. 5e), and the level of ceramide, a source of sphingosine, was significantly higher in the *TIMELESS*-knockdown cells (Fig. 5f). These results suggest that TIMELESS positively regulates S1P production via the sphingolipid salvage pathway.

**Figure 5:**
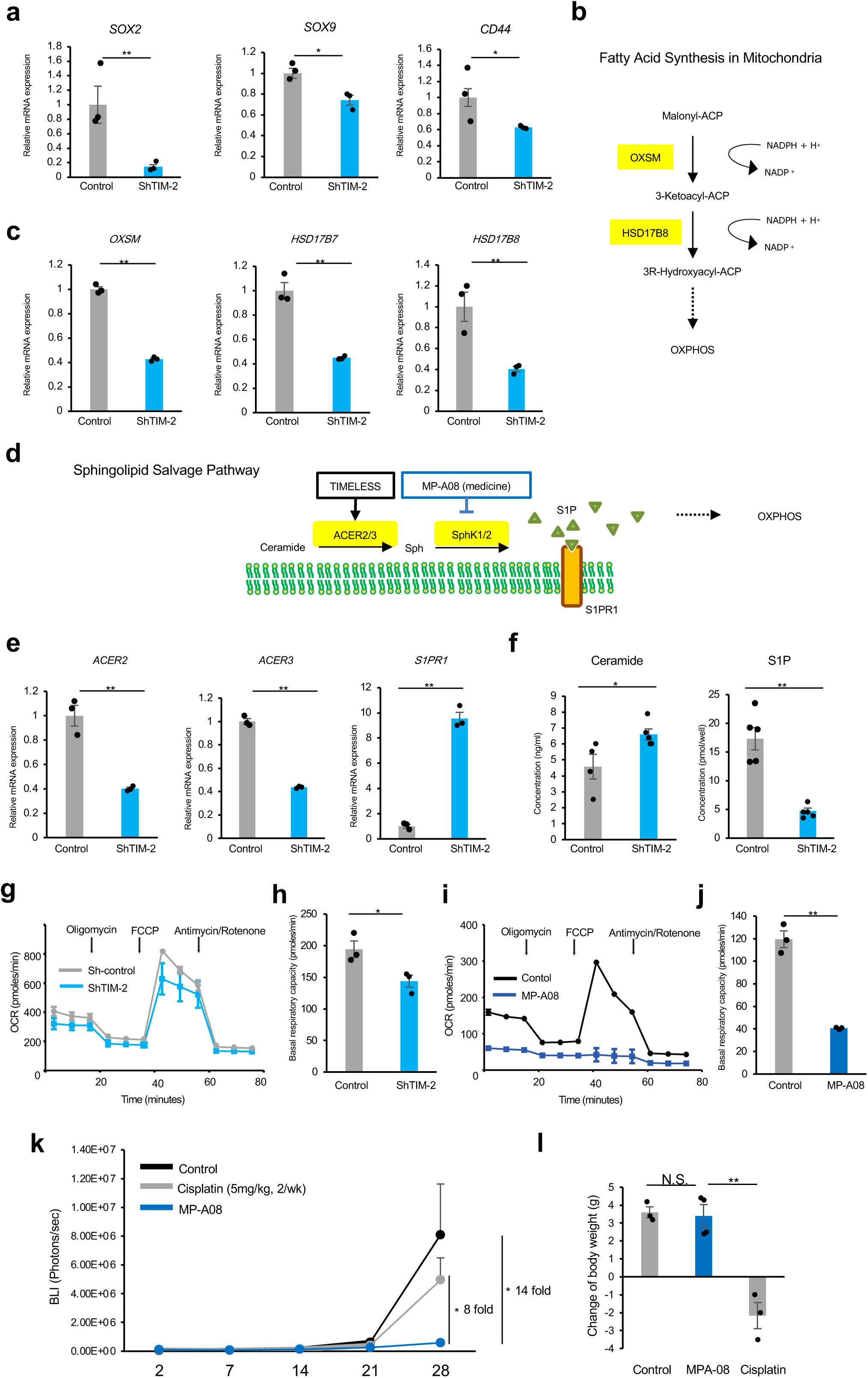
Oxidative phosphorylation regulated by TIMELESS is crucial for bone metastases. **a**, Relative mRNA expression of *Sox2, Sox9,* and *CD44* in TIMELESS-knockdown sublines and controls in PC-3 (each n=3). **b**, Schematics of fatty acid synthesis in mitochondria. **c**, Relative mRNA expression of *OXSM, HSD17B7, HSD17B8* in TIMELESS-knockdown cell line and controls in PC-3 (each n=3 independent experiments). **d**, Schematics of sphingolipid salvage pathway. **e**, Relative mRNA expression of *ACER2, ACER3,* and *S1PR1* in TIMELESS-knockdown cells and controls in PC-3 (each n=3 independent experiments). **f**, Relative concentration of Ceramide and S1P in TIMELESS-knockdown cells (each n=5 independent experiments). **g**, The OCR of TIMELESS-knockdown cells and controls (each n=3). **h,** Basal respiratory capacity of TIMELESS knockdown cells and controls in PC-3 (each n=3). **i**, The OCR of parental cells treated with MP-A08 and controls (each n=3). **j**, Basal respiratory capacity of parental cells treated with MP-A08 and controls in MDA-MB-231 (each n=5 limbs). **k**, Survival and subsequent reawakening of tumor cells in mice receiving caudal arterial injection using MDA-MB-231 (control n=8, Cisplatin n=6, MP-A08 n=8). **l**, The body weight change from Day2 to Day28 of mice in each condition (control=3, MPA-08=4, Cisplatin=3). For **a**, **c**, **e, f, h, j,** and **l**, significance was assessed by two-sided paired t-tests with the data representing means ± s.e.. *P* value for comparisons with the results are indicated (\**P*<0.05, \*\**P*<0.01). N.S., not significant.

Since two pathways, namely, fatty acid synthesis and sphingolipid salvage, are known to positively regulate oxidative phosphorylation^28,33^, we analyzed the mitochondrial metabolic state and found a decrease in the oxygen consumption rate (OCR) in *TIMELESS*-knockdown cells derived from PC-3 (Fig. 5g and 5h). The importance of the sphingolipid salvage pathway in oxidative phosphorylation was confirmed by the reduction of OCR upon treatment of MDA-MB-231 with MP-A08, an inhibitor of sphingosine phosphokinase 1/2 (Fig. 5d, 5i, and 5j). Glycolysis (represented by extracellular acidification rate, ECAR) was decreased in *TIMELESS*-knockdown cells, but unlike for OCR, MP-A08 had no effect on the basal level of ECAR (Extended data Fig. 11a–11d).

To confirm the essential role of the sphingolipid salvage pathway in bone metastasis, the effect of MP-A08 was examined in an *in vivo* metastatic model. Caudal artery injection of breast cancer MDA-MB-231 cells led to the formation of tumor masses on day 28 in NOD/SCID mice^34^, and the administration of cisplatin suppressed these masses by ∼20% (Fig. 6k). Meanwhile, MP-A08 significantly reduced the size of metastatic tumors by 93% compared with the level for untreated control cells (Fig. 5k). Notably, no body weight loss was observed in the MP-A08-administered mice, whereas cisplatin remarkably decreased the body weight of mice as an adverse effect (Fig. 5l). These results suggest the potential of MP-A08 as a future anti-metastatic drug without the adverse effect of weight loss.

## Discussion

To develop therapeutic drugs that improve the prognosis of cancer patients, there is a need to uncover the molecular mechanism behind cancer metastasis to bone. Previous studies on this issue mainly focused on the interactions between MICs and the microenvironment^8,21,25,35,36^, whereas little is known about the precise molecular machinery involved in the transcriptional regulation of reawakening and stemness. In this study, we established an MIC-reawakening model in mice and demonstrated that *TIMELESS* is a master regulator and transcription factor of the dormancy of cancer cells in bone and other organs. In contrast to the metastatic model established via the cardiac injection of cancer cells, which leads to multiple organ metastases within 2–3 months^18^, the method of injection via the caudal artery that we employed induced metastasis specifically in bone in a shorter period of time^21^. In our model, the cell number estimated using the chemiluminescence photon count plateaued within 14 days, and this latency period was attributed to the dormancy of the tumor mass^18^. The isolated MICs exhibited stem cell-like characteristics with high expression of *Sox2*, *Sox9*, and *CD44*, which may be regulated by *TIMELESS* (Extended data Fig. 11e).

Given that *TIMELESS* plays an essential role in the reawakening of MICs, the molecular machinery in MICs should be elucidated. Recently, circadian rhythm has been reported to regulate breast cancer metastasis^14^, with it being asserted that the intravasation of circulating tumor cells (CTCs) occurs during sleep in breast cancer patients^14^. The data presented in that report^14^ showed higher mRNA expression of *TRPS1* in CTCs during sleep than that during awakening. Building on the previous work, the present study involved the pseudotime analysis of single-cell ATAC-sequencing, roughly dividing MICs into early and late phases. In the early phase, chromatin accessibility of TRPS1 was specifically enhanced. The results of our analyses in combination with the previous report mentioned above^14^ suggest that TRPS1, a WNT5a-related transcription factor, might play crucial roles in the early phase in MICs, namely, in dissemination and induction of dormancy.

The late stage based on the MIC timeline was regarded as the reawakening stage. Notably, the cluster of the reawakening phase showed not only high scores for the gene accessibility of *TIMELESS*, but enhanced motif activity for *CLOCK* and *BMAL1/ARNTL. TIMELESS* has been reported as a transcriptional regulator of the circadian rhythm in *Drosophila*, while its function in mammals remains controversial^37–39^. Our results suggest the mechanisms for the upregulation of *TIMELESS* induced by or associated with *CLOCK* and *BMAL1/ARNTL* during MIC reawakening. Although *TIMELESS* is already upregulated in cancer tissue, further studies need to be performed to clarify the underlying molecular mechanisms.

In accordance with the circadian rhythm-dependent on/off machinery of *TIMELESS*, our results showed that TIMELESS protein expression levels were heterogeneous in primary cancer tissues, and the patients with high expression of TIMELESS in the primary tissue had a poor prognosis. Here, we hypothesized that cells diseeminating to bone even with low *TIMELESS* expression may be affected by the circadian rhythm mechanism and induce *TIMELESS* expression in WNT5a-enriched environments, contributing to the reawakening of MICs. Meanwhile, regarding TIMELESS-high cancer cells even in primary tumor tissues, spatial transcriptome analysis may uncover the role of *TIMELESS* in the primary tumor. Furthermore, via time-lapse monitoring focused on intravascular or post-bone metastasis, further study should be able to prospectively elucidate the dynamics of *TIMELESS* expression combined with cell cycle analysis during each metastatic process.

In this study, we showed that the sphingolipid salvage pathway is upregulated downstream of *TIMELESS*. Indeed, it has been reported that the sphingolipid pathway-related gene alkaline ceramidase 2 was regulated by transcriptional heterodimer of *TIMELESS* and transcription factor Sp1^20^. Intriguingly, Sp1 also binds to mammalian *HSD17B7* related to *HSD17B8*^40^, which has been identified as a poor prognostic factor in MICs. Given that the present study reveals that *TIMELESS* positively regulates the factors involved in mitochondrial fatty acid synthesis that lead to OXPHOS activity, *TIMELESS* regulates OXPHOS through both the sphingolipid salvage pathway and mitochondrial fatty acid synthesis. Gene enrichment analysis revealed that OXPHOS was enriched in MICs, and *TIMELESS* regulates MIC survival and reawakening through this signal.

From a clinical perspective, *TIMELESS* may be a tool for diagnosing cancer metastasis to bone. The highly sensitive detection of *TIMELESS* expression in disseminated tumor cells combined with CTCs and primary cancer tissues may enable the prediction of MIC reawakening. To date, no therapeutic strategies have been established for bone micrometastasis, but our study has clarified that the sphingolipid pathway inhibitor MP-A08 suppressed the MICs necessary for bone metastasis. Thus, the development of therapeutic agents based on MP-A08 may lead the way towards curative approaches for bone metastases. It is also possible that opaganib, a first-in-class orally administered selective inhibitor of sphingosine kinase-2, currently under a clinical trial (NCT04207255)^41^, can be applied to patients to prevent metastatic cancer recurrence.

Here, we demonstrated that *TIMELESS*, a circadian clock regulator, facilitates tumor cell survival and subsequent reawakening in bone. Clinically, *TIMELESS* and its downstream signaling pathways are potential therapeutic targets for diminishing the potency of MICs.

## Acknowledgements

We thank Dr. Masamichi Imajo (WPI-ICReDD, Hokkaido University, Japan) for providing a lentivirus plasmid vector, pLKO.1; Dr. Ryuji Matsumoto (Faculty of Medicine, Hokkaido University, Japan) for supplying cell lines (UM-UC-3 and PC-3), Ms. Mieko Hishikawa for immunohistochemistry. This study used infrastructure and equipment at GI-CoRE: GSQ in Hokkaido University. We thank Edanz (https://jp.edanz.com/ac) for editing a draft of this manuscript.

## Author contributions

H. Shimizu designed the study and performed data analysis, and H. Shimizu. and L.W. performed *in vitro* and *in vivo* experiments, all of which were supervised by L.W. and S.T. Y.S. instructed metabolic analyses. R.M. and K.K. performed single-cell data analysis. M.T., H.H., R.H., H.S. K.W. arranged immunohistochemistry, M.M. for frozen sample analysis of murine bone tissue. H. Shimizu wrote the manuscript and illustrated the figures, which was supervised and edited by N.I. and S.T. and M.T. R.A. joined the discussion, and S.T. directed the entire study.

## Material and Methods

### Cell lines

The human breast cancer luminal subtype A cell lines MCF7 (International Cell Line Authentication Committee (ICLAC) accession number CVCL_0031), the human breast cancer triple negative cell line, MDA-MB-231 (CVCL_0062), the human bladder cancer cell line UM-UC-3 (CVCL_1783), and the human prostate cancer cell line, PC-3(CVCL_0035), were cultured in Dulbecco’s modified Eagle’s medium (DMEM) containing 10% fetal bovine serum (FBS). All ethical issues relating human pathological specimens were discussed and approved by the Ethics Committee of Hokkaido University’s Graduate School of Medicine (number 022-0118)

### Establishment of the cell lines stably expressing tdTomato-Luc2

The cell lines were stably transfected with pSCII-CMV-tdTomato-Luc2 (kindly provided by Dr. Kyoko Hida, Hokkaido University), followed by selection with 0.8 mg/mL Zeocin (Invitrogen) to MCF-7; 0.4 mg/mL to MDA-MB-231; 0.5 mg/mL to UM-UC-3; 0.1 mg/mL to PC-3. The resulting live cells were sorted by fluorescence-activated cell sorting (FACS; AriaII, Becton Dickinson, Franklin Lakes, NJ) and the cells with excessive tdTomato expression were isolated. Luciferase activity was examined with a luciferase assay (Promega, Madison, WI).

### Establishment of Knockdown cell lines

The short hairpin RNAs (shRNA) targeting human TIMELESS sequences were as follows: sh-1,5’-GCTAGAGATTGTCTCCCTTAT-3’^28^, sh-2, 5’-GTGTTTGGTATTAAACATGTA-3’. The lentiviral vector, pLKO.1, was used and the transfection steps were performed according to the instructed protocol. The transfected cells were selected with 5.0 µg/mL puromycin to MCF-7: 5.0 µg/mL to UM-UC-3: 2.0 µg/mL to PC-3: 2.0 µg/mL MDAMD-231. All cell lines had been already transfected with pSCII-CMV-tdTomato-Luc2. The efficiency of knockdown was confirmed by qRT-PCR using a part of the colony.

### Animal Experiments

Balb/cA Jcl nu/nu female mice and NOD/SCID mice (CLEA Japan) were used in each animal experiment. Bioluminescent imaging (BLI) was performed with the IVIS Spectrum imaging system (Caliper Life-Sciences, Hopkinton, MA) using a post-intraperitoneal injection of Vivo Glo Luciferin In Vivo Grade (Promega). Counts of chemiluminescence were expressed as relative light units per second after injection. BLI of tumor bearing mice were acquired with IVIS Spectrum at 10 min after intraperitoneal injection of D-luciferin (50 mg/kg). The luminescence was detected under the following conditions: filter sets (emission filter=open, filter position=1 and excitation filter=block), exposure time=60s, binning=medium; and F/stop=1. For region-of-interest measurements, photon flux (photon counts per second) was measured with a setting to cover each knee of mice. All images were set on the same scale (field of view=22.2). All animal experiments were conducted in accordance with the guidelines of Hokkaido University Manual for Implementing Graduate School of Medicine and approved by the Institutional Animal Care and Use Committee at the Hokkaido University Graduate School of Medicine (number 17-0061, 22-0809). All procedures using live animal were pre-approved by the Animal Welfare Committee of Hokkaido University based on their completion of issues related to animal experiments. All ethical issues relating to animal experiments were discussed and approved by the Ethics Committee of the Hokkaido University Graduate School of Medicine (number17-0061, 22-0809).

### Establishment of *in vivo* MIC reawakening models in bone

The *in vivo* MIC reawakening models in bone was created by reference to the previous report^21^. Seven-to-eight-week-old female nude mice were used. After the mice were anesthetized with 2.00% isoflurane, the tdTomato positive cells: 1.0 *×* 10^6^ MCF-7 cells and 1.0 *×* 10^6^ of UM-UC-3 cells were injected through the intra-caudal artery. The dissemination of cancer cells was confirmed at Day0 or Day3 after the injection. The MICs were collected at fourteen days after the caudal injection. Bone-tropic tumor cells (BoTCs) were collected from bone around knee at more than 21 days after injection.

### Xenograft models using the TIMELESS knockdown sublines

Seven-to-eight-week-old female nude mice were used for the injection of 1.0 *×* 10^6^ MCF-7, 5.0*×* 10^5^ MDA-MB-231, and 1.0 *×* 10^6^ of UM-UC-3 cells. Five-to-seven-week-old NOD-SCID mice were used for 1.0 *×* 10^6^ PC-3 cells. The cells with TIMELESS knockdown and the control were prepared and injected through the intra-caudal artery and monitored by IVIS.

### Xenograft models upon the MP-A08 treatment

Eight-to-nine-week-old NOD-SCID mice were used for the injection of 5.0 *×* 10^5^ MDA-MB-231 cells. They were injected through the intra-caudal artery and monitored by IVIS. MP-A08 treatment was initiated at day 3 and MP-A08 (100 mg/kg) or vehicle (70%[vol/vol] PEG 400) was administered by intraperitoneal (IP) injection 6 times a week.

### Establishment of MICs and BoTCs cell lines from mice

MICs and BoTCs from mice were established as previously reported^18^. Briefly, bone around knee of mice were placed in a mortar filled with 3 ml of ice-cold PBS supplemented with 2% FBS and 1 mM EDTA and then crushed. Suspended cells were filtered through a cell strainer (70 µm), concentrated by centrifugation for 7 min at1,200 rpm, and resuspended in single-cell condition.

### Frozen bone sections

Frozen bone sections (8 µm thickness) were prepared as previously reported^42^. Briefly, bone samples in embedding medium (SCEM) were frozen quickly using hexane dry-ice. After trimming the frozen sample blocks, the adhesive films (Cryofilm) were mounted onto the surface. The sample were cut using the tungsten blade with 8 µm thickness. The tumor cells were verified under a fluorescence microscope to detect tdTomato.

### RNA extraction and quantitative real-time PCR

Total RNA isolation was conducted using the RNeasy Mini kit (Qiagen, Valencia, CA). cDNA was synthesized using Superscript VILO (Invitrogen, Carlsbad, CA), and Quantitative real-time polymerase chain reaction (qRT-PCR) was conducted using SYBR Green PCR Master Mix (Qiagen) and performed on the StepOne real-time PCR system (Applied Biosystems, Foster City, CA). The primer sequences are listed in Supplementary Table 1. Data were normalized to the *glyceraldehyde 3-phosphate dehydrogenase* (*GAPDH*) expression level and are expressed as fold change relative to control.

### Western blotting

Western blotting was performed according to the previous study^43^. The antibody used was as follows: TIMELESS (ab109512; abcam, Cambridge, USA).

### Immunostaining of TIMELESS in MCF7 cell block

The knockdown sublines and controls of MCF7 cells with or without CoCl_2_ treatment were fixed in formalin and cell blocks were constructed. The antibody described above was used for immunostaining of TIMELESS.

### Analysis of TIMELESS expression in human breast cancer tissues

The estrogen receptor positive, stage 3 breast cancer tissues from 212 patients who underwent surgery from 2012 to 2017 and had follow-up more than 5 years were selected. For pair-matched analyses, samples of breast cancer and bone metastasis obtained by surgery or biopsy were prepared from 26 patients. Samples of breast cancer were collected from 2005 to 2022 and those of bone metastasis were from 2015 to 2022. These clinical samples were collected from Hokkaido Cancer Center (Sapporo, Japan) and Saitama Cancer Center (Saitama, Japan). The anti-TIMELESS antibody used was described above. All ethical issues using human samples were discussed and approved by the Ethics Committee of the Hokkaido University Graduate School of Medicine, Sapporo, Japan (022-0118) and Saitama Cancer Center, Saitama, Japan (1524).

#### CoCl_2_ assay for the parent cells, MICs, BoTC3

This assay was performed as previously reported Cells (1 *×* 10^5^) suspended in 6ml of DMEM were seeded in 6-well plate, and treated with 300 µM CoCl_2_ for 10 days (from day 1 to day 11). Medium was changed every other day.

### CoCl_2_ assay for TIMELESS-knockdown sublines

The knockdown sublines of TIMELESS and the controls were seeded at a density of 1 *×* 10^5^ cells per well in 6 well plate. The medium was changed to 8 ml DMEM with CoCl_2_ treatment at the indicated days. During the CoCl_2_ treatments, no medium change was conducted.

### ELISA assay

Quantification of S1P was performed using a sphingosine 1-phosphatete ELISA Kit (K-1900; Echelon Biosciences, Salt Lake City, USA). Quantification of ceramide was done using a human ceramide kinase (CERK) ELISA Kit (MyBioSource, San Diego, USA)

### Measurement of cellular basal OCR and ECAR

The cellular metabolic assay was performed as previously reported^44^. Briefly, consumption rate (OCR) was measured using an XFp extracellular flux analyzer (Agilent Technologies, Santa Clara, CA, USA). Tumor cells were suspended in XF Assay Medium supplemented with 10 mM glucose, 1 mM pyruvate, and 2 mM glutamine. Cells were seeded in a culture plate pre-coated with Cell-Tak (Fisher Scientific). The plate was centrifuged and left to equilibrate for 60 min in a CO_2_-free incubator before being transferred to the XFp extracellular flux analyzer. The Mito Stress Test was performed as follows: 1) basal respiration was measured in XF Base Medium (1 mM sodium pyruvate, 10 mM glucose, 2 mM glutamine); 2) oligomycin (1 μM) was injected to measure respiration linked to ATP production; 3) the uncoupler carbonyl cyanide 4-(trifluoromethoxy) phenylhydrazone (FCCP, 1 μM) was added to measure maximal respiration; and 4) rotenone and antimycin A (0.5 μM each) were applied together to measure non-mitochondrial respiration. Measurement of fuel dependency and fuel flexibility was performed according to the manufacturer’s Mito fuel flex test kit protocol (Agilent Technologies).

### Microarray analysis

Total RNA was extracted from parental UM-UC-3 and its sublines (MICs and BoTC3), and microarray analysis was performed using GeneChip Expression Array (Human Clariom S Array, Affymetrix^®^).

### Screening by a cox proportional hazards model

Standardizing the transcriptomic data sets (GSE 2034 and GSE 2603) and univariate analysis (Kaplan-Mire and log rank test) were performed on the software Subio Platform ver.1.19^45^. Estrogen negative samples in the data sets were used as the control. The groups with low and high gene expression were classified, and the threshold of each gene was defined according to the distribution of histogram of the expression. P<0.05 was considered statistically significant. Cox proportional hazards models was performed by JMP^®^ version 14(SAS Institute, Inc., Cary, NC), the target genes with the significant association were selected based on the univariate analysis.

### Biological enrichment analysis

To assess the pathway enrichment, pre-ranked version of gene sets enrichment analysis (GSEA) and DAVID Functional Annotation Clustering Tool were used. Gene sets derived from the Kyoto Encyclopedia of Genes and Genomes (KEGG) pathway database was also used to visualize the pathway.

### Transcriptomic data analysis of clinical primary and metastatic samples

Using the software Subio Platform ver.1.19, the following procedures were conducted: Standardizing the transcriptomic data set deposited in GitHub and principal component analysis. Wilcoxon signed-rank test was performed on JMP^®^ version 14 (SAS Institute, Inc., Cary, NC). P<0.05 was considered statistically significant.

### ATAC-seq library preparation

Single-cell ATAC-seq libraries were prepared using a SureCell ATAC-Seq Library Prep Kit (Bio-Rad) and a SureCell ddSEQ Index Kit (Bio-Rad) according to the manufacturer’s instructions. Libraries were loaded at 1.5 pM on a NextSeq 550 (Illumina) and sequencing was performed using the following read protocol: Read 1: 118 cycles; i7 index read: 8 cycles; and Read 2: 40 cycles. FASTQ files were processed using the ATAC-Seq Analysis Toolkit (Bio-Rad) to generate debarcoded and aligned read data.

### scATAC-seq analysis—ArchR

ArchR v.1.0.2 for scATAC-seq analysis^30^ was applied. All analyses were performed with the hg19 genome assembly using ArchR’s “addArchRGenome (“hg19”)” function. ATAC-seq quality was evaluated using by enrichment of ATAC-seq accessibility at transcription start sites. ArchR’s “createArrowFiles ()” function calculated TSS enrichment score and unique nuclear fragments. The cells were filtered using cut-off of TSS enrichment score of eight and 3000 unique fragments per cell to exclude low-quality cells. To filter out doublets, ArchR’s “addDoubletScores ()” with “k = 10, knnMethod = “UMAP”, LSIMethod =1” parameters was used. Quality plots were made by “plotGroups ()”, “plotFragmentSizes ()”, and “plotTSSEnrichment ()”.

ArchR’s “addIterativeLSI ()” function with genome-wide 500-bp tile matrix was used for calculating iterative LSI information. The cells were clustered by ArchR’s “addClusters ()” with Seurat’s “FindClusters ()” and default parameters. To run uniform manifold approximation and projection (UMAP), ArchR’s “addUMAP ()” function with 40 nearest neighbors were used.

Gene activity scores were visualized in the UMAP overlay by ArchR’s “plotEmbedding ()” function. A gene score matrix was obtained using “getMatrixFromProject ()” for getting pre-imputed matrix, “imputeMatrix ()” for matrix imputation with MAGIC, and log2 (Imputed gene score + 1) for normalization. Psudotime for each cell was obtained by slingshot pseudotime.

TF activities were measured in ArchR: ChromVAR deviation scores. For ChromVAR analysis, the raw insertion counts for all peaks were used as input. HOMER motif annotations were added by ArchR’s “addMotifAnnotations ()” function. the GC bias-corrected motif deviation scores were computed using ArchR’s “addDeviationsMatrix ()”. The motif score matrix was obtained as same as the gene score matrix described above. Psudotime for each cell was obtained by slingshot pseudotime.

### Statistical analysis

Graphical data are presented as the mean and standard deviation (S.E.) and Student’s t test and log rank test were used to compare variables. *P*<0.05 was considered statistically significant.

---

**Supplemental table 1.**
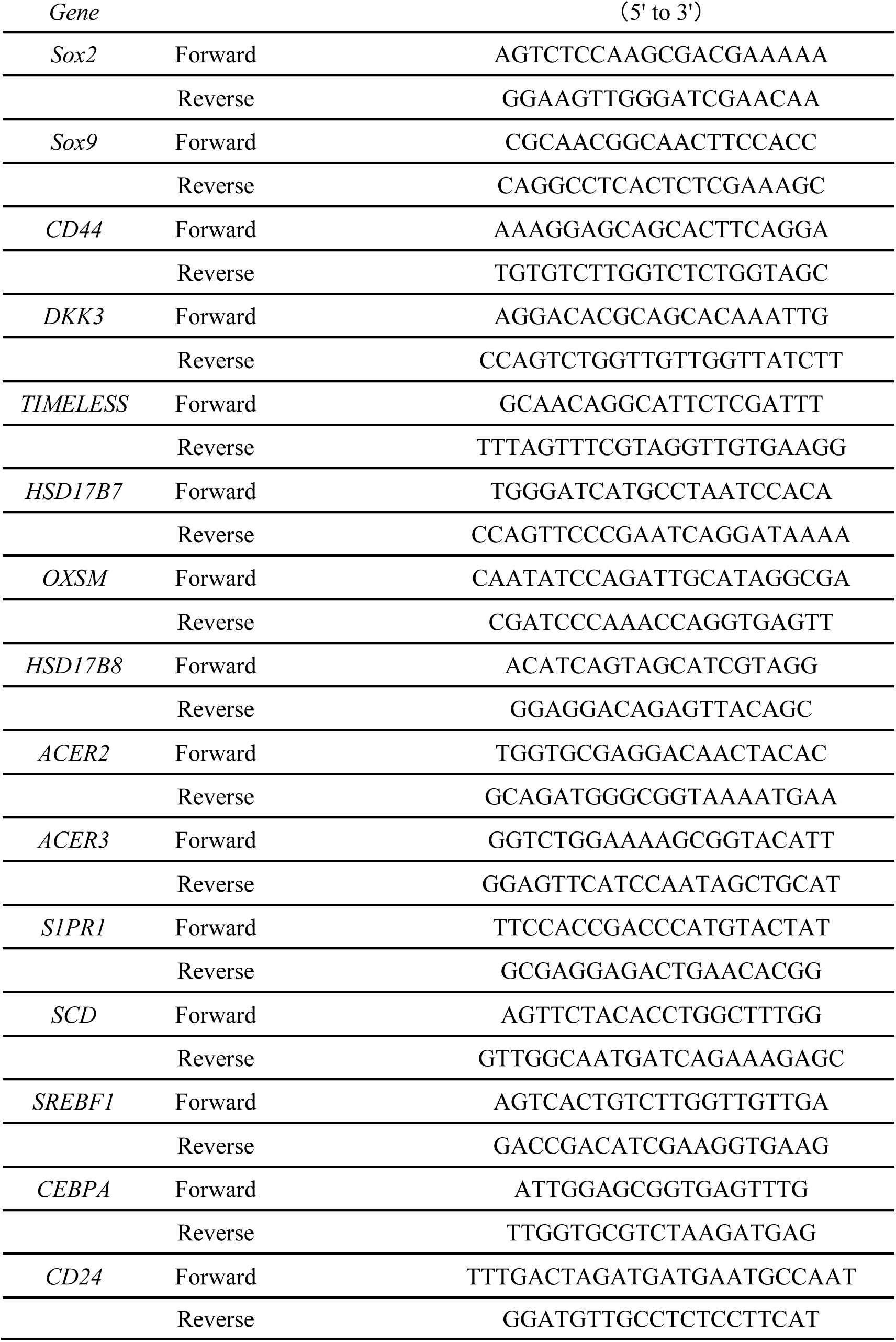
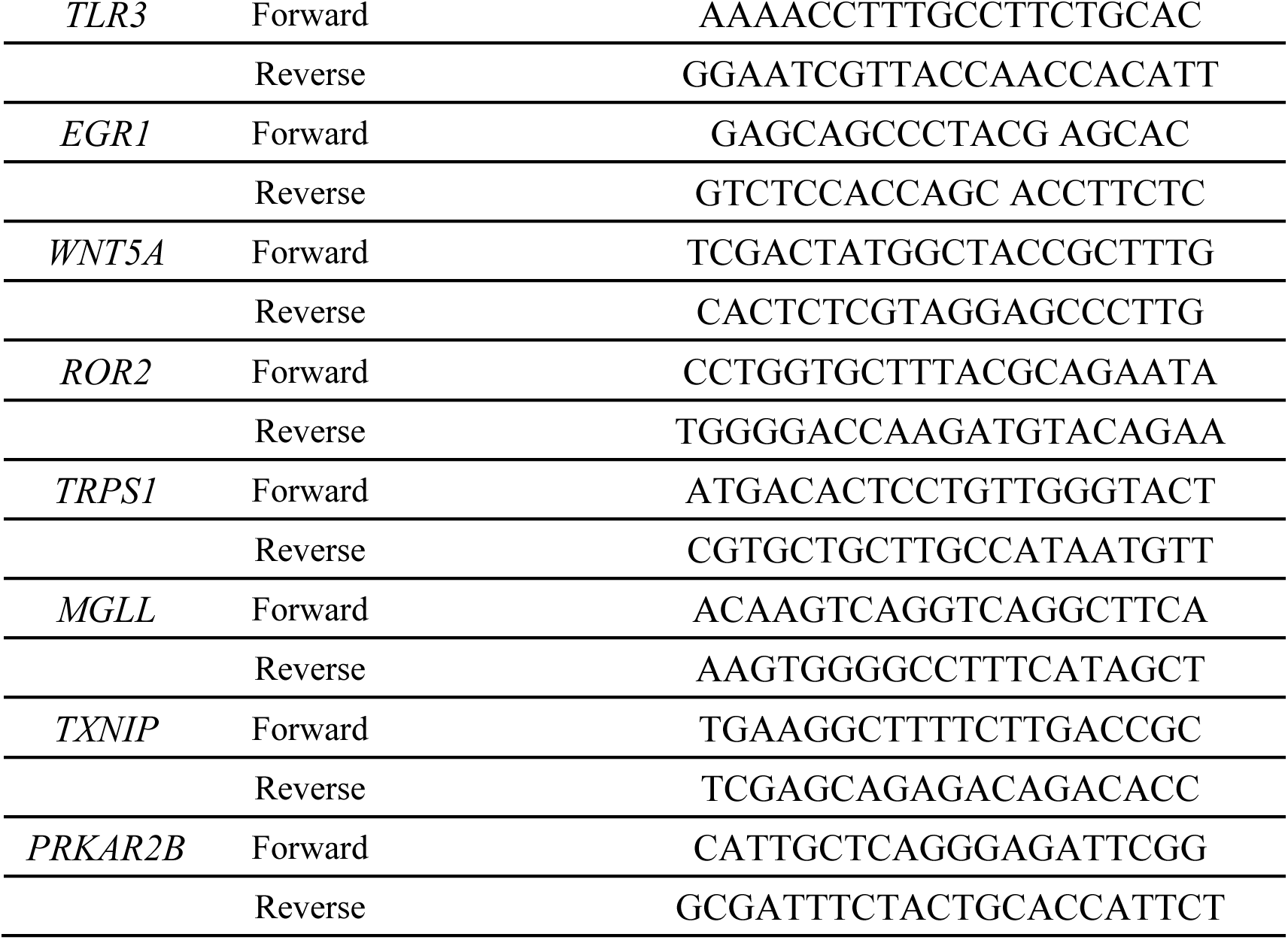

**Extended data Fig. 1.**
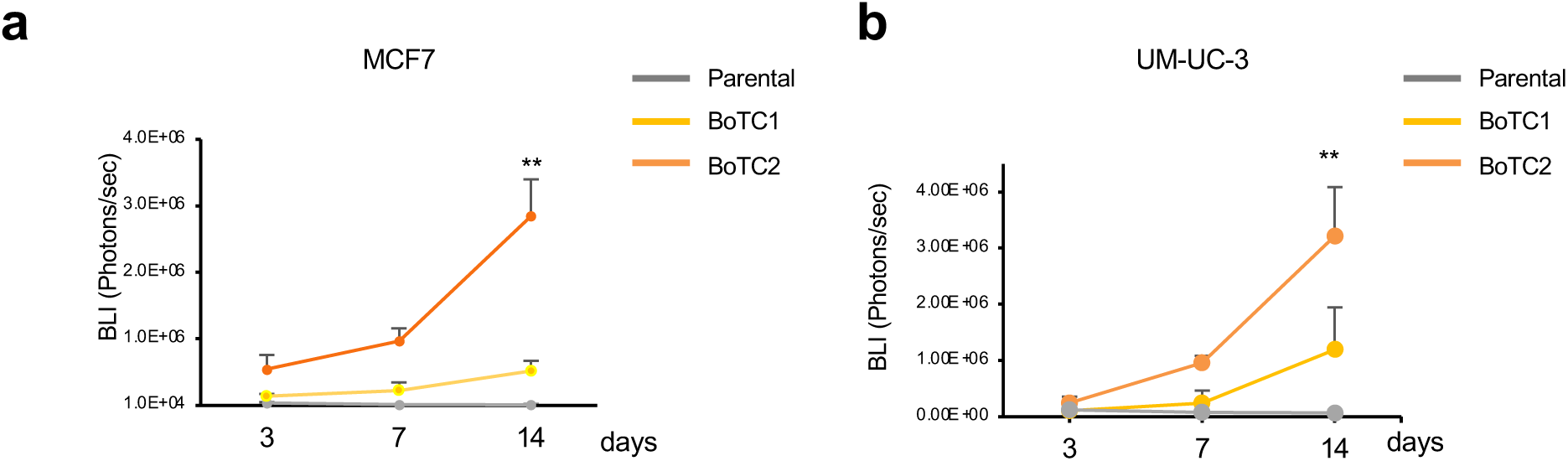
The signal intensities during the latency period. **a, b,** Quantification of BLI signal in bone metastatic lesion, BoTC1 and BoTC2 of MCF7 (**g**) and UM-UC-3 (**g**) cells (n=6 limbs with metastatic lesion in each parental cell line and BoTC1 of MCF7, and n=8 limbs with metastatic lesion in BoTC1 of UM-UC3, BoTC2 of MCF7 and UM-UC3). For **a, b,** significance was assessed by two-sided paired t-tests with the data representing means ± s.e.. *P* value for comparisons with the results are indicated (\**P*<0.05, \*\**P*<0.01).

**Extended data Fig. 2.**
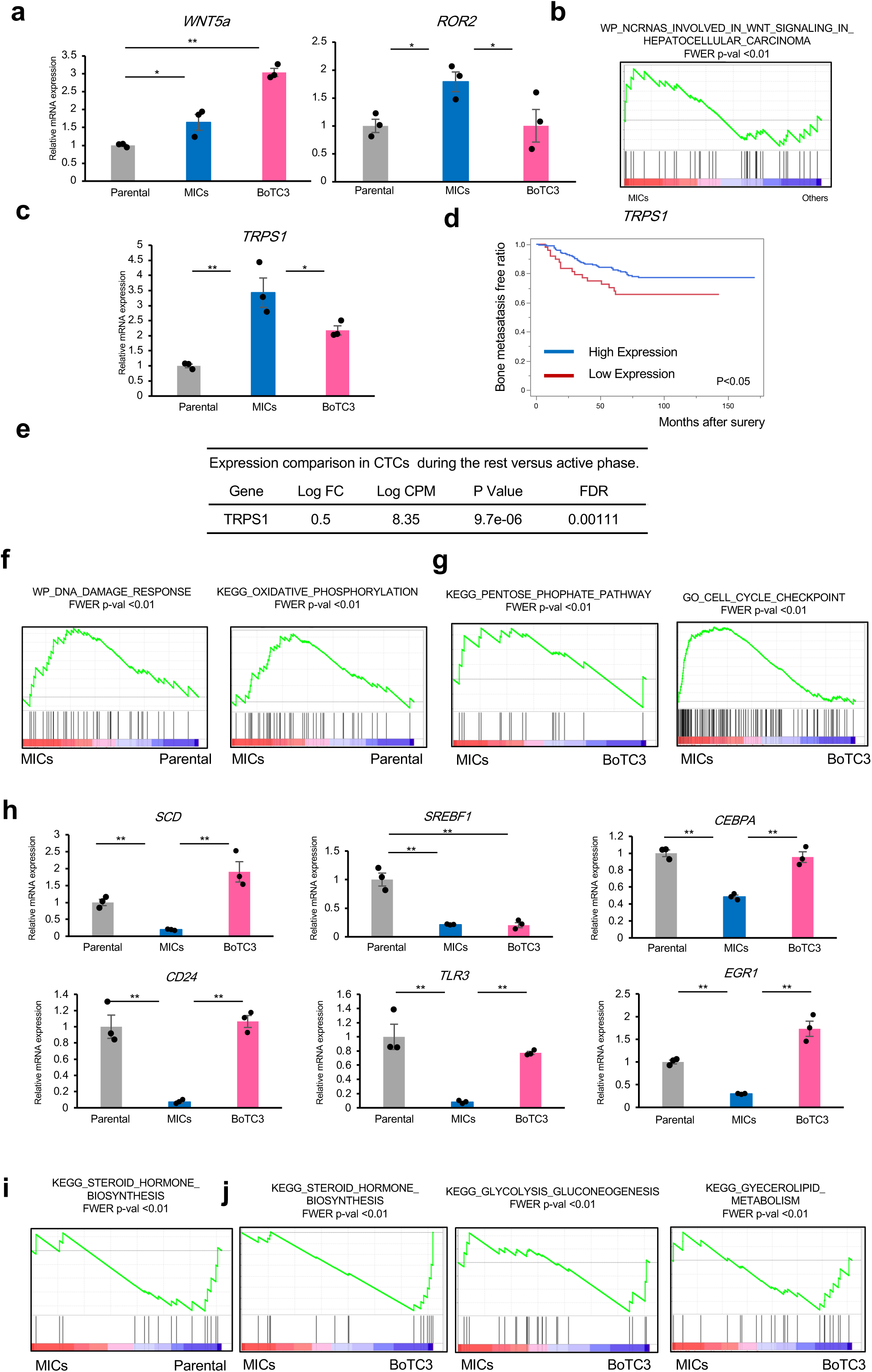
Profiles of MICs. **a**, Relative mRNA expression of *WNT5a, ROR2,* in parental cells, MICs, and BoTC3 (each n=3 independent experiments). **b**, Gene-expression signature that associate with WNT signal phosphorylation in MICs compared with the others. **c**, Relative mRNA expression of *TRPS1* in parental cells, MICs and BoTC3. **d**, Proportion of bone metastasis free patients (from the MSKCC/EMC cohort primary tumor dataset) stratified according to *TRPS1*. Statistical analysis was performed by log ranked test **e**, Relative mRNA expression of TRPS1 in CTCs during rest and active phase (from the GSE180097). **f**, Gene-expression signatures that associate with DNA damage response (left) and oxidative phosphorylation (right) in MICs compared with parental cells. **g**, Gene-expression signatures that associate with metabolism (left) and cell cycle checkpoint (right) in MICs compared with BoTC3. **h**, Relative mRNA expression of *SCD, SREBF1, CEBPA, CD24, TLR3,* and *EGR1* in parental cells, MICs and BoTC3. (each n=3 independent experiments.) **i**, Gene-expression signature that associate with steroid hormone biosynthesis in MICs compared with parental cells. **j**, Gene-expression signatures that associate with metabolism in MICs compared with BoTC3. steroid hormone biosynthesis (left), glycolysis gluconeogenesis (middle), and gyecerolipid metabolism (right). For **a, c, e, h,** significance was assessed by two-sided paired t-tests with the data representing means ± s.e., and for **d**, significance was assessed by log ranked test. *P* value for comparisons with the results are indicated (\**P*<0.05, \*\**P*<0.01).

**Extended data Fig. 3.**
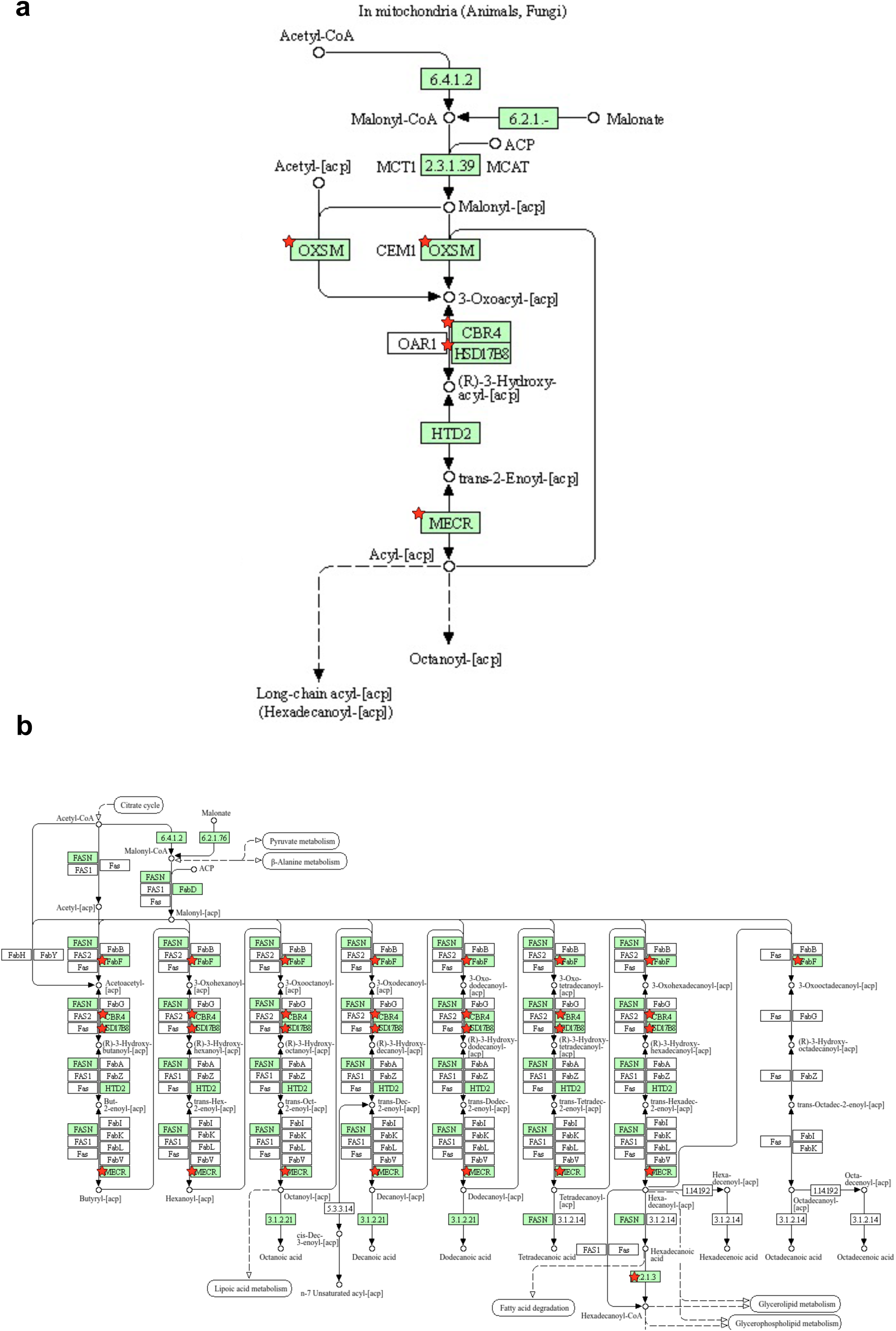
Upregulated metabolic pathway in MICs. **a**, Mitochondrial fatty acid synthesis as an upregulated pathway in MICs compared with parental cells. **b**, Sphingolipid pathway as an enriched pathway in MICs compared with parental cells.

**Extended data Fig. 4.**
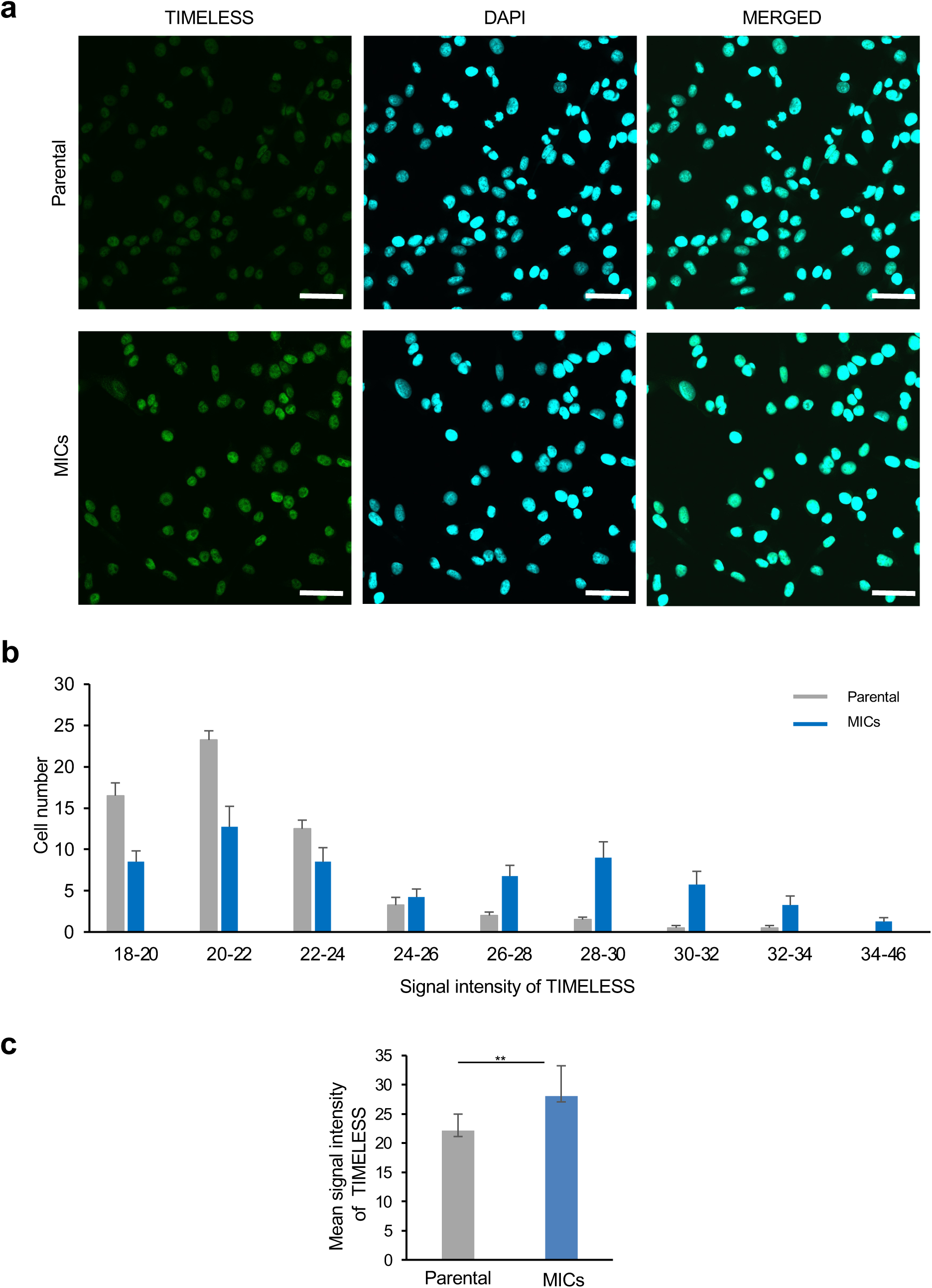
Bimodal expression of TIMELESS in MICs. **a**, Representative immunofluorescence images of TIMELESS in parental cells and MICs. Nuclei were stained by DAPI. Scale bar: 100 µm. **b**, Histogram of signal intensity of TIMELESS immunofluorescence in parental cells and MICs (60 cells were randomly extracted and the intensity was calculated. The operation were repeated 5 times.) **c**, Mean signal intensity of TIMELESS immunofluorescence in parental cells and MICs was displayed. For **c**, significance was assessed by two-sided paired t-tests with the data representing means ± s.e.. *P* value for comparisons with the results are indicated (\**P*<0.05, \*\**P*<0.01).

**Extended data Fig. 5.**
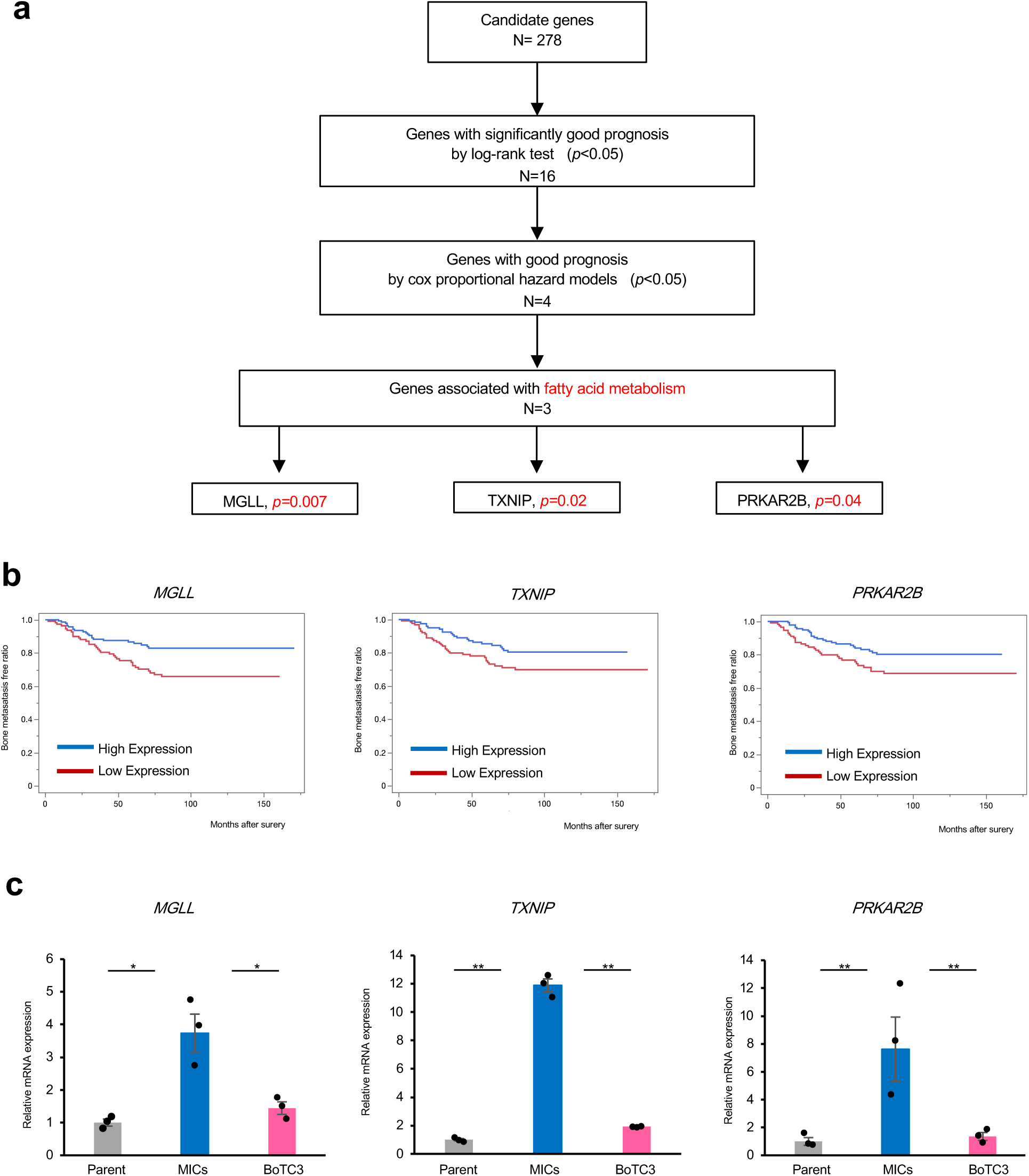
Screening with clinical transcriptome datasets identifies the independent favorable prognostic factor. **a**, The independent favoroble prognostic genes in upregulated genes in MICs, extracted by a screening model with the MSKCC/EMC cohort primary tumor dataset. **b**, Proportion of bone metastasis free patients (from the MSKCC/EMC cohort primary tumor dataset) stratified according to *MGLL*, *TXNIP*, and *PRKAR2B*. **c**, Relative mRNA expression of *MGLL*, *TXNIP*, and *PRKAR2B* in parental cells, MICs and BoTC3 (each n=3 independent experiments). Significance was assessed by two-sided paired t-tests with the data representing means ± s.e.. *P* value for comparisons with the results are indicated (\**P*<0.05, \*\**P*<0.01).

**Extended data Fig. 6.**
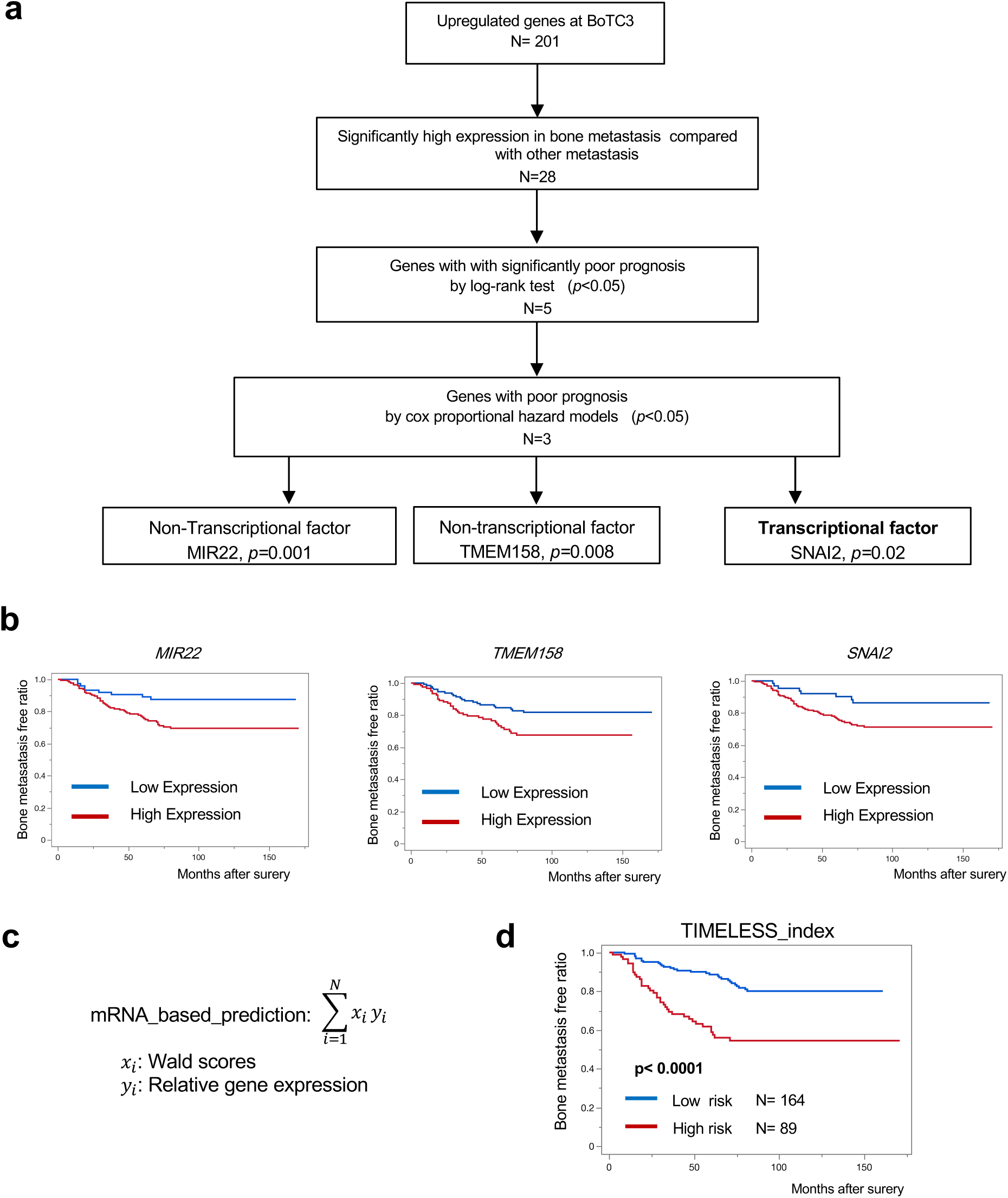
Screening with clinical transcriptome datasets identifies poor prognostic factors in BoTC3. **a**, The independent poor prognostic genes in upregulated genes in BoTC3, extracted by a screening model with the MSKCC/EMC cohort primary tumor dataset. **b**, Proportion of bone metastasis free patients (from the MSKCC/EMC cohort primary tumor dataset) stratified according to *MIR22*, *TMEM158*, and *SNAI2*. **c**, Prognosis prediction formula based on the 11 extracted genes by the clinical transcriptome screen. **d**, Proportion of bone metastasis free patients based on the prognosis prediction formula from the MSKCC/EMC cohort primary tumor dataset (low risk, n=164; high risk, n=89) Significance was assessed by log ranked test.

**Extended data Fig. 7.**
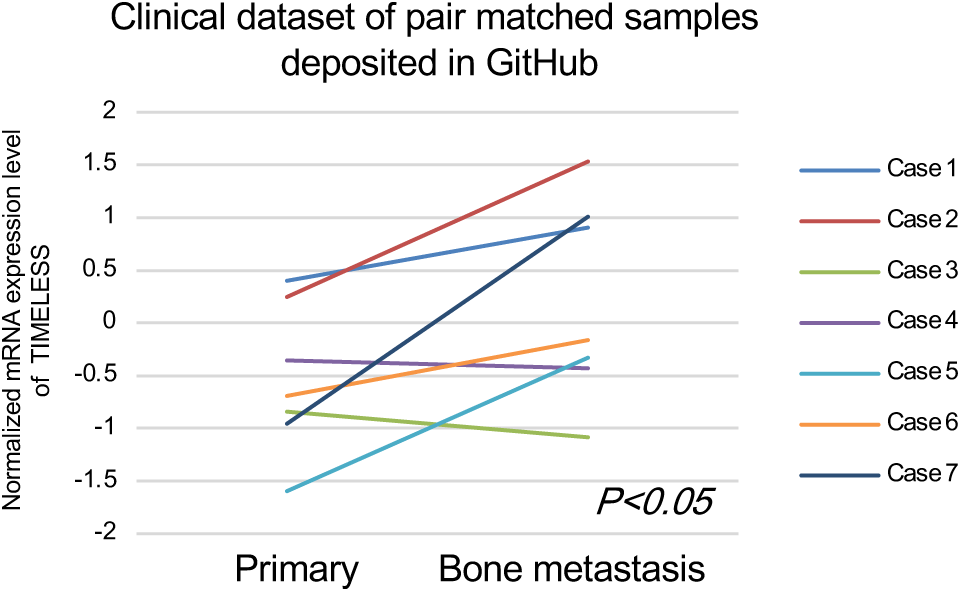
Pair-mathed analysis using public data. Pair-matched analysis of normalized mRNA expression of *TIMELESS* on ER^+^ stage 3 or 4 breast cancer primary and matched bone metastasis tumor (n=7; from Github jciInsight_2017). Pairs are connected with a line and Wilcoxon signed-rank P value is shown.

**Extended data Fig. 8.**
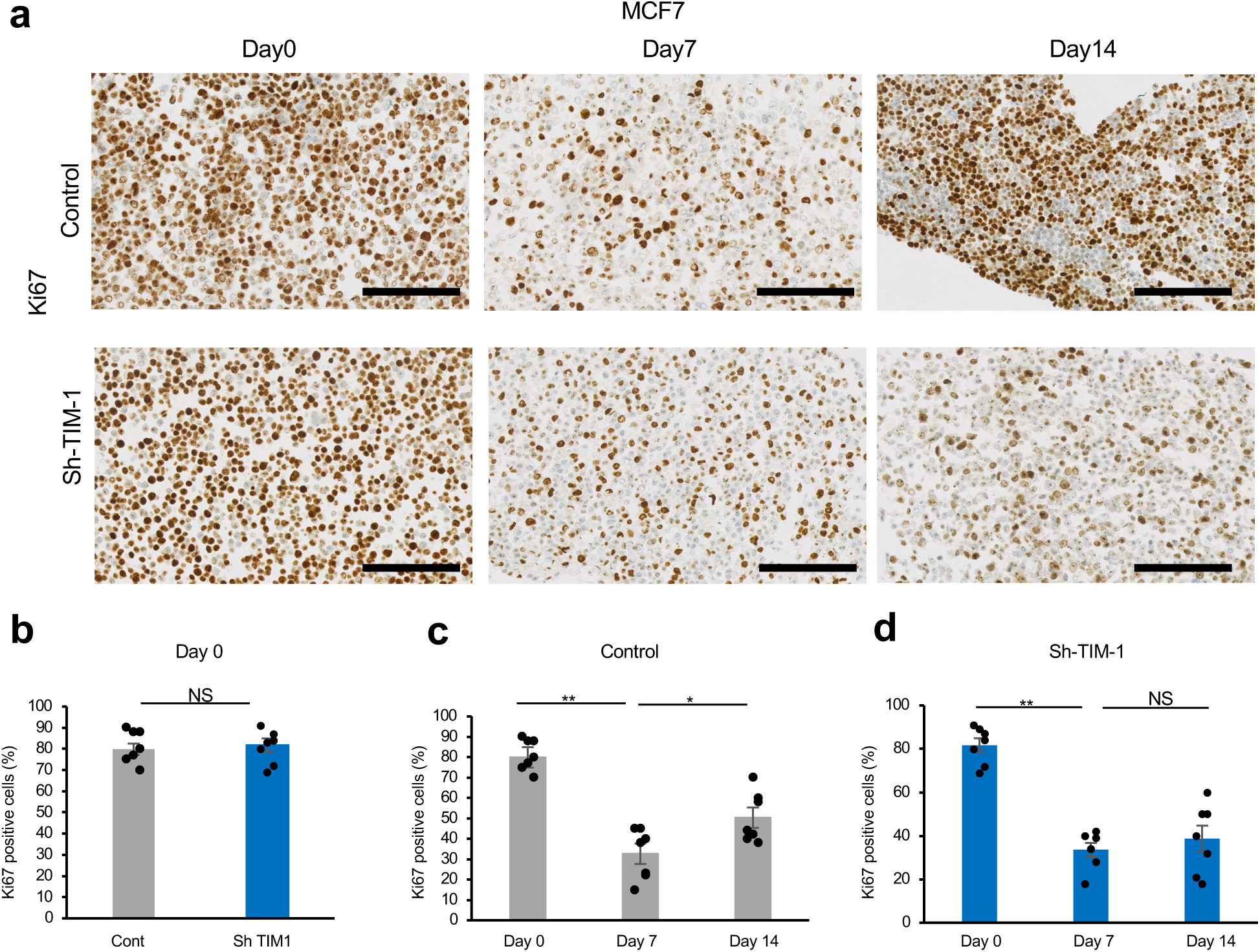
Serial changes in Ki67 of TIMELESS knock down cell lines under CoCl2 treatment. **a**, Representative Ki67 immunostaining images of TIMELESS-knockdown cell lines and controls of MCF7 under CoCl_2_ condition (Day0, before CoCl_2_ treatment; Day 8, during CoCl_2_ treatment; Day 14 after CoCl_2_ treatment). Scale bar: 200 µm. **b**, Positive ratio of Ki67 in TIMELESS-knockdown cell lines and controls of MCF7 cells at Day 0. **c**,**d**, Positive ratio of Ki67 in controls (**c**) and TIMELESS-knockdown (**d**) of MCF7 cells (Day0, before CoCl_2_ treatment; Day 8, during CoCl_2_ treatment; Day14, after CoCl_2_ treatment) (each n=7 independent experiments) For **b** and **d**, significance was assessed by two-sided paired t-tests with the data representing means ± s.d.. *P* value for comparisons with the results are indicated (\**P*<0.05, \*\**P*<0.01). NS, not significant. ECRA: extracellular acidification rate.

**Extended data Fig. 9.**
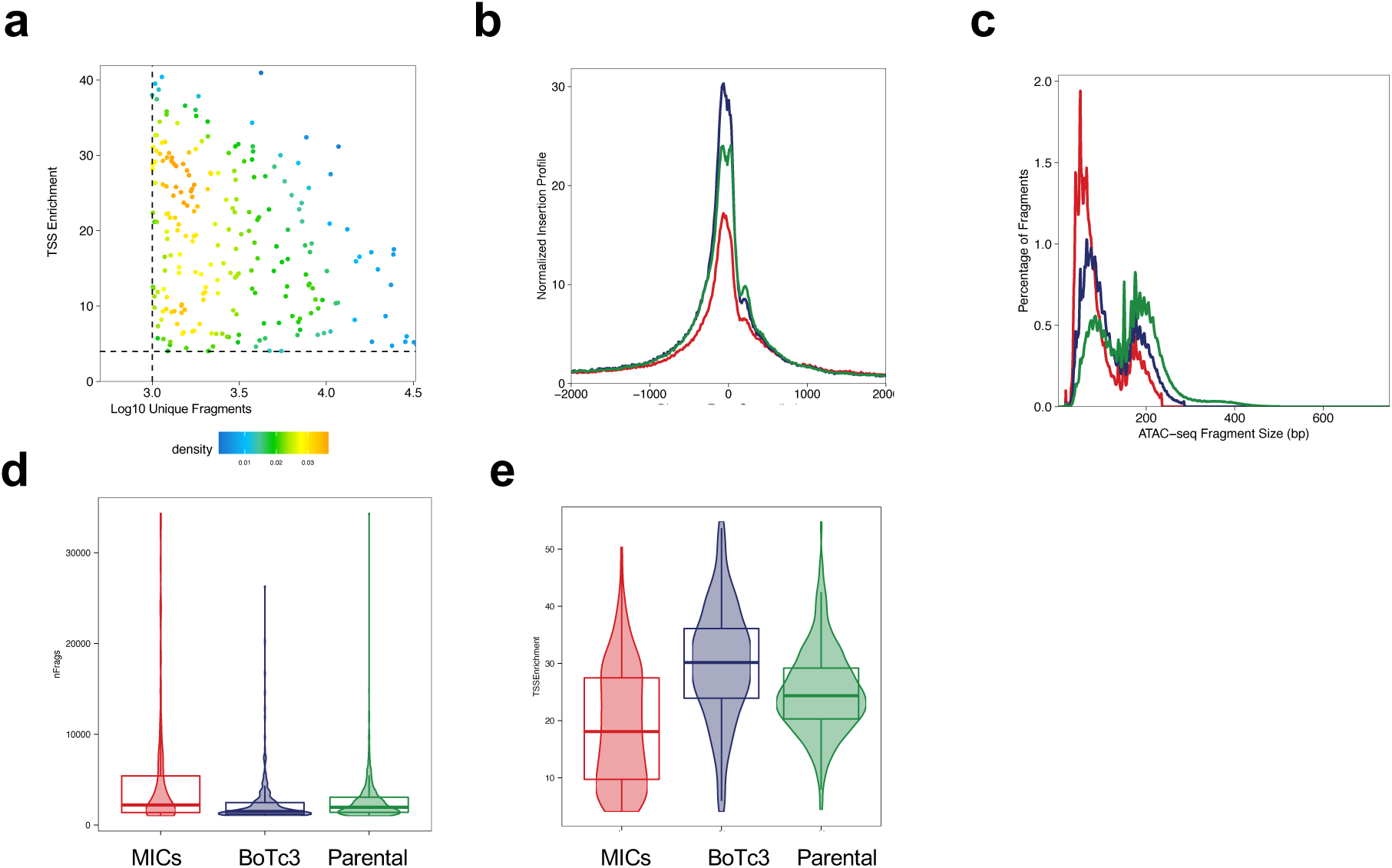
scATAC-sq quality control. **a**, Scatter plots of TSS enrichment score and number of fragments. Each dot represents each MIC. Cells shown in the figure were defined as high quality and further analyzed. **b-e**, scATAC-seq quality control. TSS enrichment in **b** and **d**, insertion profiles in **c**, number of fragments in **e** for each subline red; MICs, blue: BoTC3, green: Parental.

**Extended data Fig. 10.**
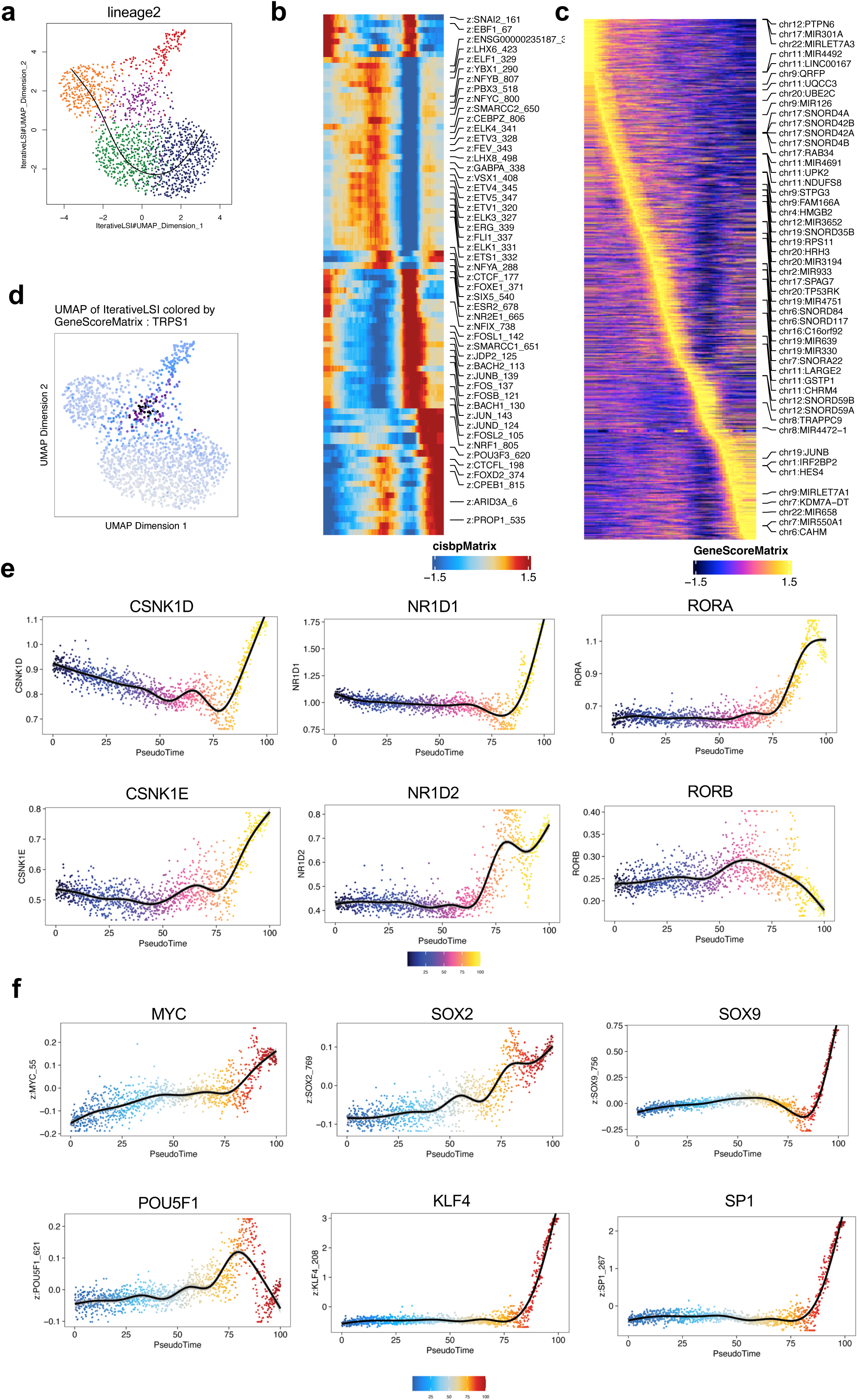
Epigenetic land scapes on MICs and BoTC3. **a**, Pseudotime trajectory analysis starting from the cluster 2 to the cluster 5. **b**, Pseudotime heatmap ordering of most variable chromVAR TF motifs across the lineage2. **c**, Pseudotime heatmap ordering of gene accessibility scores of the most variable genes across the lineage2. **d**, UMAP projection colored by gene activity scores, reflecting the general chromatin accessibility of TRPS1. **e**, Gene accessibility scores of CLOCK-controlled genes along pseudotime trajectory. **f**, Motif enrichments of stemness-related genes along the psuedotime trajectory.

**Extended data Fig. 11.**
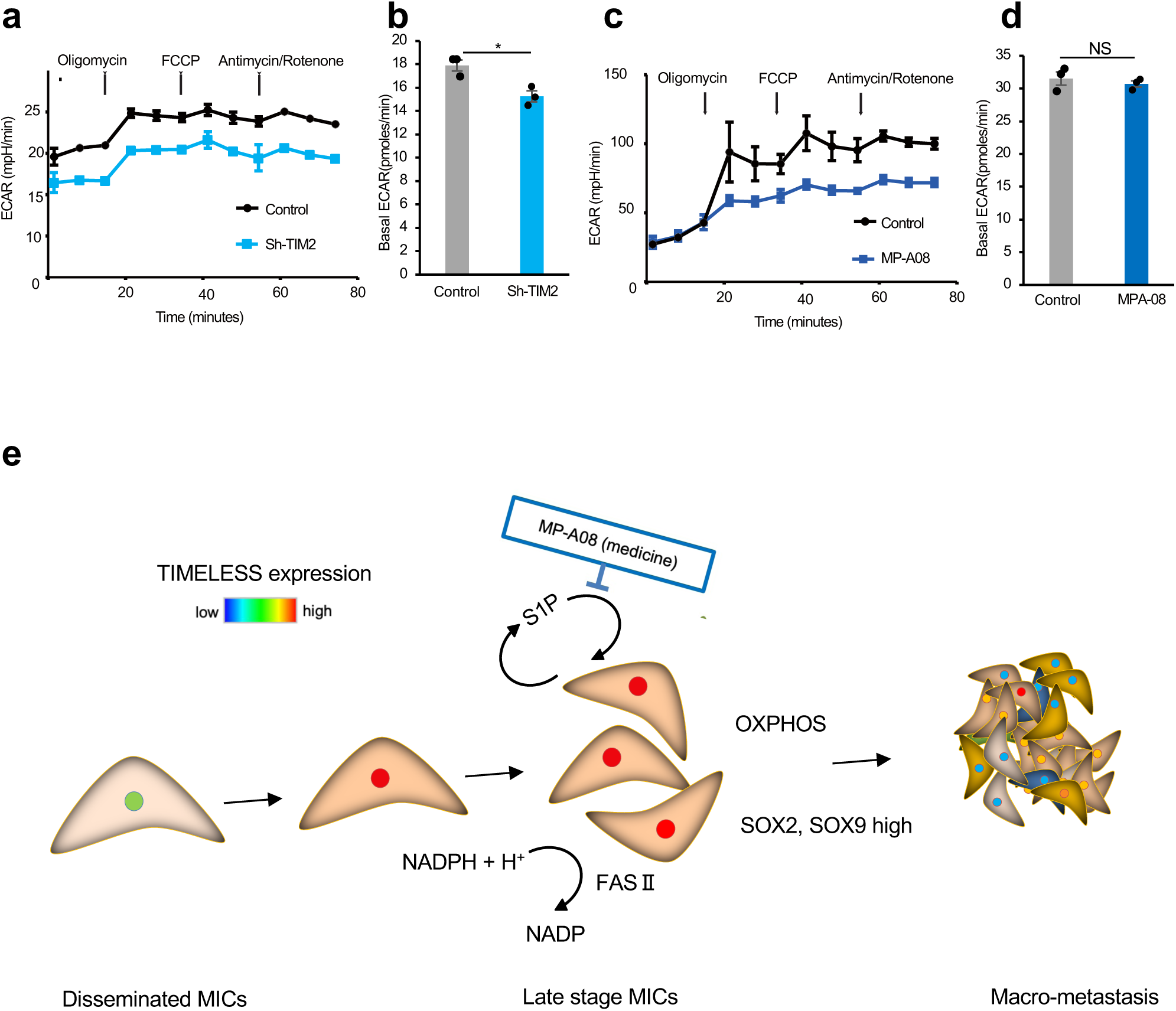
Metabolic profiles of TIMELESS knock down cells. **a**, ECAR of TIMELESS-knockdown cell lines and controls of PC-3. **b**, Basal ECAR of TIMELESS-knockdown cell lines and controls of PC-3 (n=3). **c**, ECAR of parental cells treated with MP-A08 and controls in PC-3. **d,** Basal ECAR of parental cells treated with MP-A08 and controls in PC-3 (n=3). **e**, Schematic illustration of survival and reawakening expression of TIMELESS in MICs and subsequent macro-metastasis. For **b** and **d**, significance was assessed by two-sided paired t-tests with the data representing means ± s.e.. *P* value for comparisons with the results are indicated (\**P*<0.05, \*\**P*<0.01). NS, not significant. ECRA: extracellular acidification rate.

